# Development and Validation of Thalia: A High-Resolution Pediatric Computational Model of a 10-Month-Old Infant

**DOI:** 10.64898/2026.07.09.737638

**Authors:** Alice Albrecht, Georgios Ntolkeras, Lilla Zölleia, Georgios A. Sideris, Francesca Marturano, Michael H. Lev, P. Ellen Grant, Giorgio Bonmassar

**Author notes:** Corresponding author, Email address (Giorgio Bonmassar). Contributed equally to this work (Alice Albrecht, Georgios Ntolkeras).

## Abstract

Numerical human models are essential to advance medical device design, safety assessment, and study how anatomical development influences physiological processes. Despite increasing availability of pediatric models, a critical gap remains in high-resolution, non-morphed whole-body models representing children around one year of age. Existing pediatric models are often derived from morphing older anatomies or lack sufficient tissue segmentation to accurately capture early developmental anatomy.

This study introduces Thalia, a non-morphed, high-resolution numerical model of a healthy 10-month-old female. The model was constructed by segmenting 442 tissues from Magnetic Resonance Imaging data. Brain tissues were automatically segmented using an infant-specific FreeSurfer framework, followed by semi-automated and manual refinement in 3DSlicer. The model was validated by expert review and quantitative comparison with age-matched anatomical values reported in the literature. The resulting whole-body model provides detailed anatomical representation across the brain, musculoskeletal system, vasculature, and internal organs, enabling realistic assignment of tissue-specific properties for computational studies. It provides a versatile platform for pediatric medical device development, dosimetry, safety assessment, and bioelectromagnetic simulations. This pediatric numerical anatomical model is openly available as an open-source resource.

**Highlights:**

- High-resolution model of a 10-month-old child developed from MRI data
- 442 tissues segmented to capture early anatomical development accurately
- Model validated against expert review and age-matched anatomical data
- Illustrative Example of a pediatric transcranial magnetic stimulation use case

## 1. Introduction

Computational modeling has become an indispensable tool across biomedical science and engineering, providing mechanistic insight into complex biological systems and supporting the design, optimization, and safety evaluation of emerging medical technologies. By enabling the simulation of multiscale biological interactions and device-tissue interfaces, these models allow researchers to perform controlled virtual experiments that are impractical, unethical, or infeasible to conduct in vivo. In addition, computational models support early stage in silico assessments, offering quantitative predictions of biological responses, informing design decisions, and de-risking subsequent preclinical and clinical studies.

### 1.1. Evolution of Computational Phantoms

Computational phantoms have evolved from early stylized geometric models to highly detailed, anatomically realistic representations. Initial stylized phantoms, such as the Medical Internal Radiation Dosimetry (MIRD) (Siegel et al., 1999), International Commission on Radiological Protection (ICRP) (Inter-national Commission on Radiological Protection, 1991), and Oak Ridge National Laboratory (ORNL) (Han et al., 2006) models, provided basic frameworks for dose calculation but lacked anatomical real-ism. The introduction of voxel phantoms in the 1980s, based on medical imaging such as Computed Tomography (CT) and Magnetic Resonance Imaging (MRI), significantly improved anatomical fidelity. Notable examples include the ICRP 110 Reference Phantoms (Zankl, 2010), the Visible Human Project (Ackerman, 1998), and the eXtended CArdiac-Torso (XCAT) family (Segars et al., 2015), which also incorporates physiological motion.

Since the 2000s, Boundary Representation (BREP) and hybrid phantoms have enabled flexible, anatomically detailed, and computationally efficient modeling using techniques such as Non-Uniform Rational B-Splines (NURBS) and polygonal meshes. Representative examples include the Golem phantom (Zankl and Wittmann, 2001), Mesh-Based Computational Phantoms (MCPs) (Carter et al., 2019), and the University of Florida hybrid reference phantoms (Lee et al., 2010). More recent models further incorporate population diversity (Chinese Family of Phantoms (Pi et al., 2017)), physiological variability (IT’IS Virtual Family (IT’IS Foundation, 2026a)), and specialized conditions such as pregnancy (Maynard et al., 2014; Kopacin et al., 2022), thereby supporting accurate dose assessment across diverse clinical and research contexts. In addition, recent efforts have focused on the development of pediatric head-only models, such as the Neonate Head Model project (Song et al., 2013), which provide detailed anatomical representations for neuroimaging and bioelectromagnetic applications. However, these models are limited to the head and do not provide whole-body anatomical coverage.

### 1.2. Advances and Gaps in Pediatric Computational Phantoms

Pediatric models reflect age-related anatomical variations (Table 1), but notable limitations remain. For example, the Nina model, representing a 3-year-old female, was created by morphing the Roberta model, which represents a 5-year-old (Gosselin et al., 2014), a process that can introduce anatomical inaccuracies. Similarly, the Charlie model, which represents an eight-week-old infant from the GSF Family (Petoussi-Henss et al., 2002), includes only 45 segmented tissues and has undergone limited validation.

**Table 1:** Summary of Pediatric Computational Phantoms below 5yo.

| Model Name | Model Age | Released Year | Imaging Used | Tissues Number | Morphing Used |
| --- | --- | --- | --- | --- | --- |
| UF Family of reference hybrid phantoms (Lee et al., 2010) | 1-year-old, 4-year-old, 5-year-old | 2010 | CT | 47 | Yes |
| Charlie (GSF Family, Baby) (Petoussi-Henss et al., 2002) | 8-week-old | 2014 | CT | 56 | No |
| Nina (ViP Family) (IT'IS Foundation, 2026a) | 3-year-old | 2014 | Roberta | 75 | Yes |
| Roberta (ViP family) (IT'IS Foundation, 2026a) | 5-year-old | 2014 | MRI | 78 | No |
| Pediatric xCAT phantoms (Segars et al., 2015) | Newborn, 1-year-old, 2-year-old, 5-year-old | 2015 | PET-CT | 64 | Yes |
| Chinese Family phantoms (Pi et al., 2017) | 5-year-old | 2017 | RPI mesh phantom | 50 | Yes |
| Martin (ViP family) (Jeong et al., 2021) | 29-month-old (2.5-year-old) | 2021 | MRI-CT | 114 | No |

To address these limitations, the Martin model (a 29-month-old male) (Jeong et al., 2021) and the Athena model (a 3.5-year-old female) (Ntolkeras et al., 2023) were developed to provide detailed, anatomically accurate representations of toddlers, overcoming the limitations associated with morphing approaches. However, highly detailed models for younger infants around one year of age remain scarce. To fill this gap, we introduce Thalia, a 10-month-old female model based on high-resolution MRI data, offering unprecedented anatomical detail and supporting sex-specific representation in pediatric computational modeling.

### 1.3 Illustrative Example: Transcranial Magnetic Stimulation

Computational models are essential for improving treatment strategies, including presurgical mapping in epilepsy, where the epileptogenic zone, defined as the brain region responsible for seizures, must be removed while preserving surrounding functional areas. In adults, functional MRI can identify these functional zones by having patients perform tasks (Szaflarski et al., 2007). In children, particularly infants and toddlers, anesthesia is often required during MRI, making task-based functional Magnetic Resonance Imaging (fMRI) infeasible (Raschle et al., 2012). An alternative approach combines structural MRI with Transcranial Magnetic Stimulation (TMS) to map functional regions (Garvey and Mall, 2008; Narayana et al., 2024; Rajapakse and Kirton, 2013).

TMS is a non-invasive method that induces electric currents in targeted brain regions via magnetic stimulation. It can activate or inhibit neural tissue without requiring the patient to be awake, making it suitable for younger children. Different TMS coils vary in depth penetration and focality. As an illustrative application of the Thalia model, we performed low-frequency (LF) finite element method (FEM) simulations in Sim4Life using a magnetoquasi-static formulation to estimate the electric field distributions generated by multiple TMS coil designs (Deng et al., 2020). This application provides an example of how the model can be used to investigate coil-dependent stimulation characteristics in the infant brain. The simulations are intended to illustrate the utility of the model rather than to identify optimal coil designs or provide clinical recommendations for pediatric TMS.

### 2. Methods

### 2.1. Subject and Data Acquisition

A 10-month-old female subject was selected based on the quality and completeness of the available MRI sequences, representative age-appropriate body metrics, and the absence of anatomical abnormalities (Figure 1 and Table 2), as well as the availability of multiple imaging sequences that facilitated the segmentation process. Two experienced neuroradiologists, P.E.G. and M.H.L., each with more than 20 years of clinical experience, evaluated the images to confirm their suitability for segmentation (Figure 2). The subject’s body metrics were representative of typical children of her age (Body Mass Index (BMI), 17.8 kg/m2, 55th percentile at 8 months of age, close to the age of acquisition of the MRI images (Centers for Disease Control and Prevention, 2024)), ensuring the relevance of the model for pediatric research applications. The images were sourced from the Boston Children’s Hospital (BCH) Picture Archiving and Communication System (PACS) database. The study protocol was approved by the BCH Institutional Review Board, with written informed consent waived due to the retrospective nature of the study, in compliance with the Health Insurance Portability and Accountability Act (HIPAA).

**Table 2:** MRI sequence parameters used for segmentation.

| Sequence Name | Resolution (mm) | TR(ms)/TE(ms)/TI(ms)/FA(°) | FOV (mm) | NSA |
| --- | --- | --- | --- | --- |
| COR IR (full body) | 1.25 x 1.25 x 5.0 | 3380/53/220(TI)/120 | 756x338 | 2 |
| COR T1 (full body) | 0.73 x 0.73 x 5.0 | 624/9.4/140 | 799x309 | 1 |
| SAG IR (spine) | 0.52 x 0.52 x 3.0 | 2990/111/200(TI)/120 | 358x202 | 1 |
| AX IR (neck, chest) | 0.94 x 0.94 x 5.0 | 4988/261/200(TI)/120 | 240x300 | 2 |
| AX T2 HASTE FS (neck, chest) | 0.80 x 0.80 x 3.5 | 1600/97/160 | 222x280 | 1 |
| AX T2 FLAIR FS (fast)(brain) | 0.39 x 0.39 x 3.0 | 9000/85/2500(TI)/150 | 200x162 | 1 |
| SAG T1 MPRAGE (fast)(brain) | 0.94 x 0.94 x 1.0 | 1680/2.43/958(TI)/9 | 210x210 | 1 |
| COR T1 MPRAGE (fast)(brain) | 0.94 x 0.94 x 1.0 | 1680/2.43/958(TI)/9 | 198x148 | 1 |
| AX T1 MPRAGE (fast)(brain) | 0.94 x 0.94 x 1.0 | 1680/2.43/958(TI)/9 | 192x165 | 1 |

**Figure 1:**
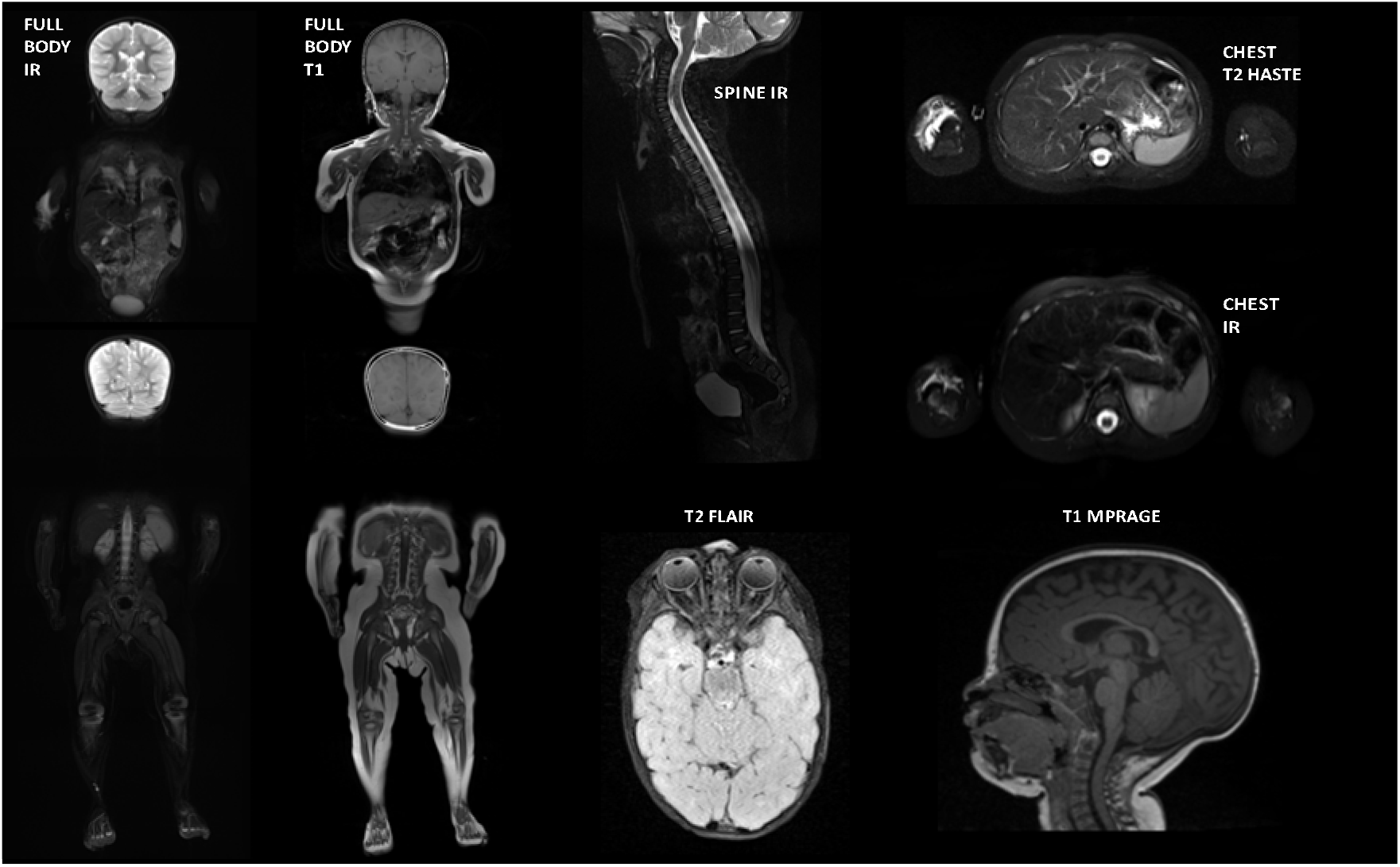
Structural MRI scans used for the segmentation process. Different imaging scans were employed to visualize body tissues with varying levels of detail. T1w MPRAGE was used for segmenting brain structures, while T2w FLAIR was utilized to segment the brain’s arteries and veins. The spine was delineated using spine IR imaging. The segmentation of other tissues was achieved through a combination of full-body T1 and IR images, along with T2 HASTE and IR chest scans.

**Figure 2:**
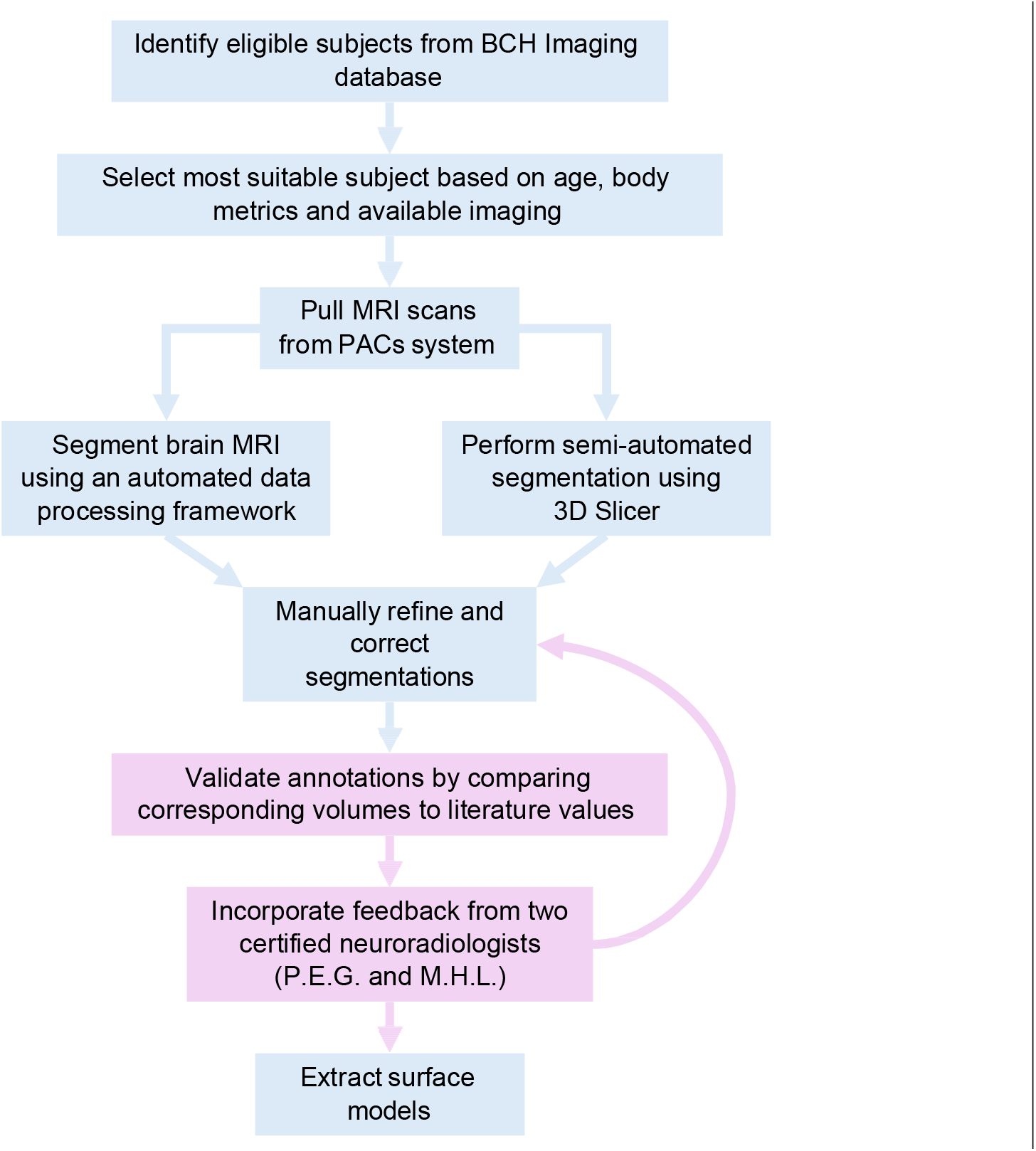
Flowchart of the segmentation. The first step was to select a female patient with available whole-body sequences, with anatomy and body metrics representative of the general population for her age and sex. We then pulled all the available MRI scans from the BCH PACS database. The process of segmenting different body tissues was different for brain and non-brain tissues. For brain structures segmentation, an automated infant-specific data processing framework was initially used, prior to the semi-automated and manual techniques applied for non-brain tissues. Semi-automated tools in 3D Slicer included segment subtraction and addition, merging, region growth, grow-from-seed, and smoothing. The results were reviewed by two subspecialized neuroradiologists (P.E.G. and M.H.L) and cross-referenced with the literature (boxes highlighted in pink). All tissues were compiled into the reference T1 sequence, and final tissue surfaces were extracted to generate the model.

### 2.2. Data Processing and Registration

The tissue segmentation labels were reviewed across multiple MRI sequences, including (Inversion Recovery (IR), T1, T1 Magnetization Prepared Rapid Gradient Echo (MPRAGE), T2 Fluid-Attenuated Inversion Recovery (FLAIR), T2 Half-Fourier Acquisition Single-Shot Turbo Spin Echo (HASTE)), which together covered the entire body from head to toe, including all four extremities (Figure 1 and Table 2). The medical images were resampled to a uniform spatial resolution of 0.5 × 0.5 × 0.5 mm3. Linear registration was performed between MRI sequences using the whole-body coronal T1-weighted (T1w) image as the reference volume. The registration process involved six degrees of freedom (rigid registration): rotation and translation along the x-, y-, and z-axes. The 3D Slicer General Registration (BRAINS) tool (Fedorov et al., 2012) was used to perform this co-registration.

### 2.3. Segmentation

Segmentation of the anatomical model was achieved through a combination of automated, semi-automated, and manual techniques to ensure high precision.

The initial automated segmentation of the brain was performed using an infant-specific framework within the FreeSurfer software package (Fischl, 2012). Infant Freesurfer is specifically designed to address the unique challenges of analyzing infant brain MRI data during the first 2-3 years of life, taking into account the rapid developmental changes and the substantial anatomical variability in this postnatal cohort. The volumetric segmentation component of Infant FreeSurfer (infantFS) relies on a multi-atlas label fusion approach, in which a subset of the training dataset, the age-matched nearest neighbors of the input, was used to guide the determination of the most appropriate anatomical labels at each voxel, based on both atlas information and MRI image features (Zöllei et al., 2020; FreeSurfer Development Team, 2025). A total of 32 anatomical labels were assigned to the brain MRIs. This framework provided a robust foundation for segmenting the infant brain anatomy (Figure 5a-d).

Following the automated segmentation, semi-automated and manual refinements were made to correct inaccuracies and refine the segmentation. This included addressing the underestimation of gray matter boundaries and improving the segmentation of specific structures such as the pons (Figure 3). Additional precision was achieved by segmenting fine anatomical structures like the colliculi, mammillary bodies, pituitary gland (hypophysis), and hypothalamus, as well as arteries and veins, to ensure vascular continuity throughout the model (Figure 5f). Further segmentation was carried out using 3D Slicer (Fedorov et al., 2012) for tissue labels not included in the infantFS output. This included detailed semi-automatic segmentation of the eyes and surrounding structures (lens, choroid, ciliary muscle, sclera, cornea, retina, vitreous body, extraocular muscles, and infraorbital fat). The skull segmentation (Figure 5e) was derived from the overall anatomical model, and outer tissues, including muscles, fat, and skin, were delineated. Cerebrospinal Fluid (CSF) was also identified and segmented in the remaining areas. Semi-automated tools in 3D Slicer included segment subtraction and addition, merging, region growth, grow-from-seed, and smoothing. These operations were used to improve segmentation accuracy and consistency across tissues. Importantly, surface smoothing was applied for visualization purposes only and did not alter the underlying anatomical model used for simulations. The smoothing visible in rendered figures was performed using the “surface smoothing” function in the 3D Slicer display settings, with smoothing fac-tors typically ranging from 0.1 to 0.3 depending on the anatomical structure. After finalizing the head and neck segmentation, the body was also segmented using semi-automatic tools in 3D Slicer, followed by manual refinement through an iterative feedback process involving two neuroradiologists (Figure S1). Whole-body (non–organ-specific) segmentations included full-body vessels (Figure 4c), connective tis-sues, subcutaneous fat, and skin (Figure 4a). Thoracic organs and thoracic cage segmentation comprised the lungs, heart, thymus, and associated thoracic soft tissues, as well as the complete thoracic cage, including the sternum and ribs. Abdominal organs included the liver, spleen, pancreas, stomach, small and large intestines, kidneys, adrenal glands, and gallbladder. Pelvic organs included the urinary bladder and reproductive organs, namely the uterus, vagina, ovaries, and fallopian tubes (Figure 6A). Spinal segmentation included the entire spine with all vertebrae, as well as associated neural and supporting tissues: spinal cord, cauda equina, spinal meninges, cerebrospinal fluid (CSF) of the spine, intervertebral discs, and spinal connective tissues (Figure 6B). The limb skeletal structures were segmented, comprising the complete upper and lower skeleton (Figure 4b). For each bone, Compact Bone (CB) and Bone Marrow (BM) were segmented separately to ensure accurate tissue-specific material property assignment in the computational model.

**Figure 3:**
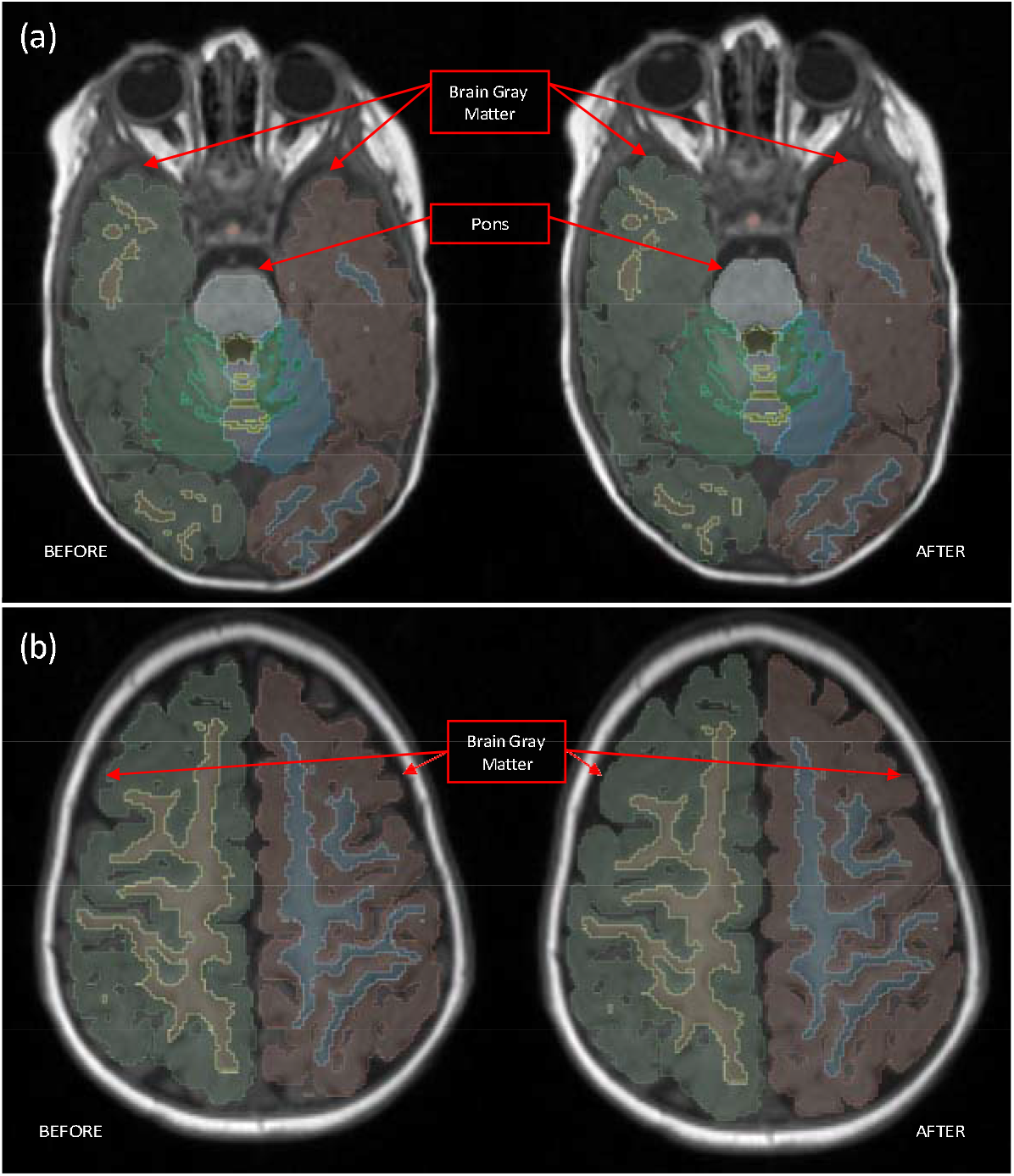
Examples of Brain Auto-Segmentation Corrections. (a) The top panel shows corrections that were applied to the automated segmentation results of the cerebral gray matter and the pons boundaries on axial view images. (b) The bottom panel illustrates corrections applied to the gray matter CSF boundary, also shown in the axial view.

**Figure 4:**
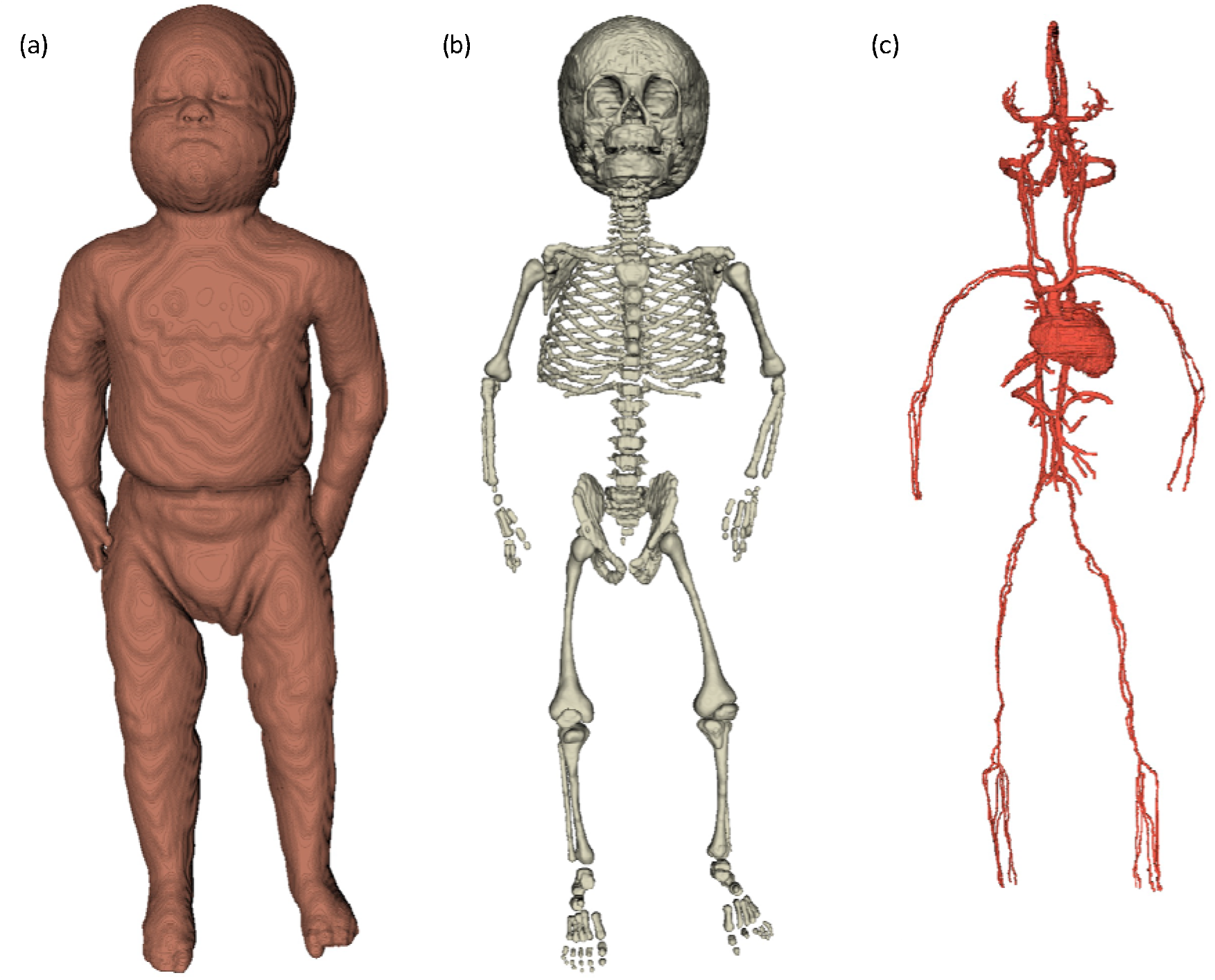
Whole-body tissue segmentation of the 10-month-old girl. (a) Whole-body volume segmentation from T1-weighted MRI. (b) Bone segmentation across the full body. (c) Vessel segmentation, including extremities, head, and cardiac cavities, based on T1 and inversion recovery images. MPRAGE and T2-FLAIR sequences were also used for brain vessel extraction.

**Figure 5:**
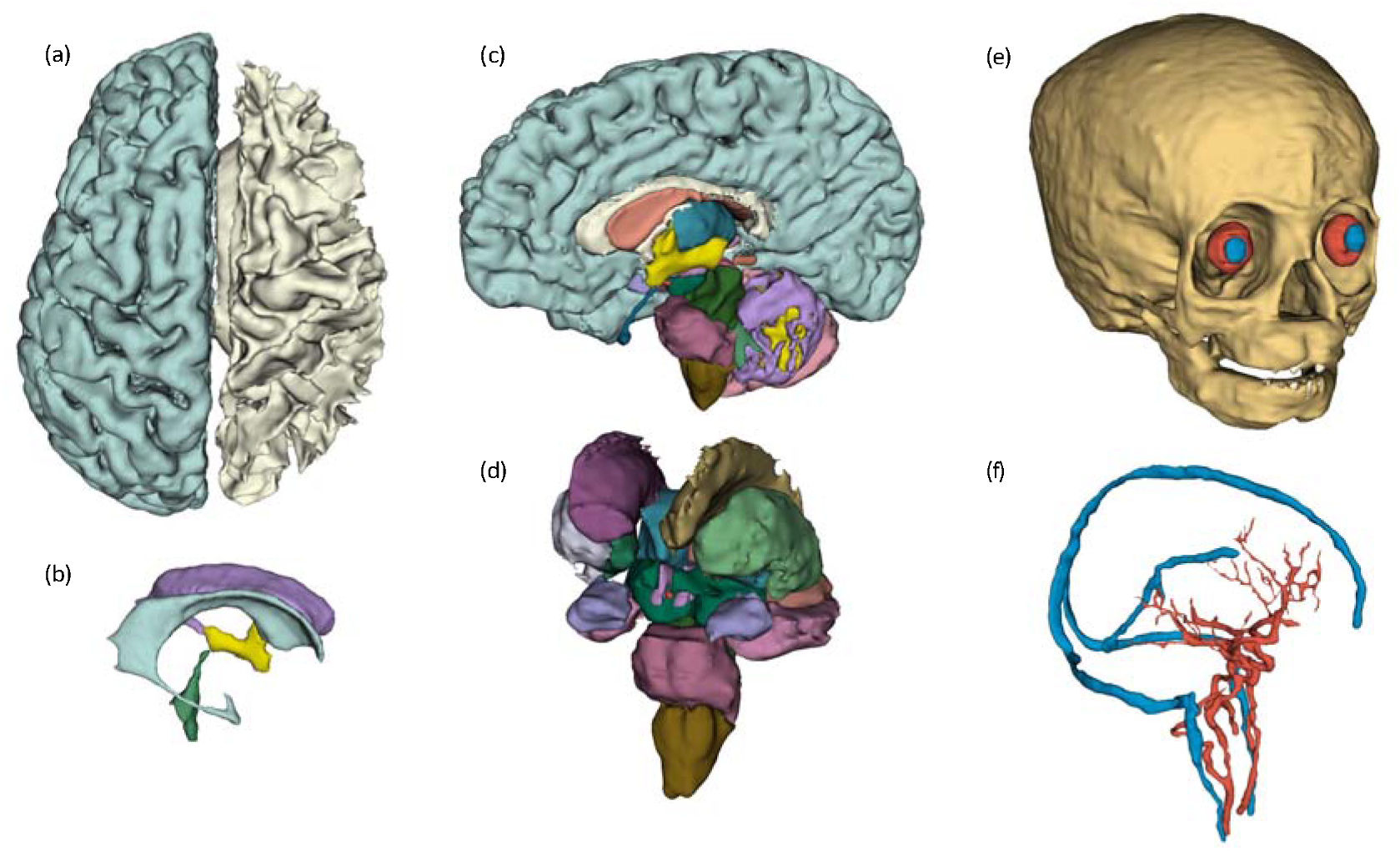
Segmentation of brain, cranial bones, and vascular system of the 10-month-old girl. **(a)** Left hemisphere gray matter and right hemisphere white matter, showing cortical gyri and sulci as well as white matter tract organization. **(b)** Ventricular system containing CSF; the left and right lateral ventricles are shown in purple and blue, the third ventricle in yellow, and the fourth ventricle in green. **(c)** Medial view of the brain revealing deeper brain regions, cerebellar structures and the brainstem. **(d)** Subcortical structures including the medulla oblongata (brown), pons (dark pink), ventral diencephalon (dark green), hypothalamus (light purple), thalamus (blue), and midbrain (green, partially obscured). **(e)** Anterior view of the skull illustrating cranial bones, orbital and nasal cavities, and dental structures. **(f)** Cerebrovascular system highlighting the 3D distribution of major cerebral vessels, with arteries in red and veins in blue.

**Figure 6:**
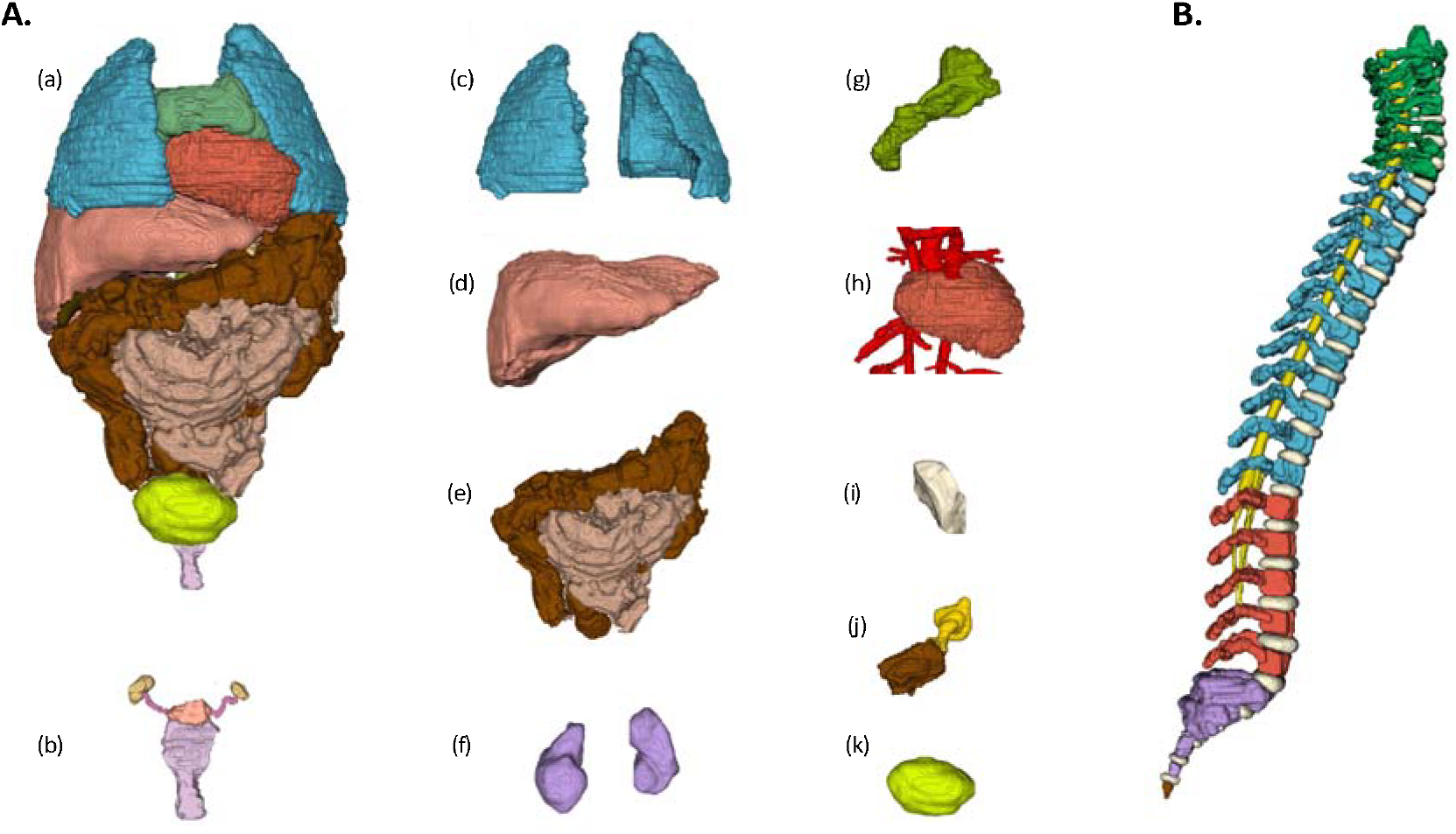
Segmentation of thoracoabdominal organs and spinal structures of the 10-month-old girl. **(A)** Thoracoabdominal organs: (a) all thoracoabdominal organs, (b) female genitalia including ovaries (yellow), fallopian tubes (pink), uterus (coral), and vagina (purple), (c) lungs, (d) liver, (e) large bowel wall (dark brown) and small bowel wall (light brown), (f) left and right kidneys, (g) pancreas, (h) heart muscle and vessels, (i) spleen, (j) stomach contents (brown) and air (yellow), (k) urinary bladder. **(B)** Spine segmentation showing the 7 cervical (green), 12 thoracic (blue), 5 lumbar (red), 5 sacral (purple) vertebrae, the coccyx (brown), intervertebral discs (white), and spinal cord (yellow). Each vertebrae was separately segmented (Supplementary Table S1). Colors in panel **(A)** and **(B)** are independently selected.

Finally, the segmented volume was imported into MATLAB for voxel reclassification that were not initially assigned to a specific tissue, using a 3D neighborhood analysis to assign the most frequently occurring neighboring label to each voxel. The final segmented tissues were exported in STL format using 3D Slicer, ensuring compatibility with Sim4Life for TMS simulations.

### 2.4. Tissue Properties

Accurate simulations require precise knowledge of tissue properties, including mass density and dielectric properties, consisting of the relative permittivity and the electrical conductivity. These properties are highly dependent on the frequency of exposure (Ibrani et al., 2012). For our simulations of TMS, we assumed an electric current of 5 kA and a sinusoidal magnetic flux density of 2.5 kHz, consistent with the frequency of the biphasic pulse used in commercial TMS systems (Crowther et al., 2013). This waveform represents an idealized approximation of TMS excitation and was not calibrated to a specific stimulator (e.g., in terms of dB/dt). All coils were driven using this same waveform, enabling a controlled comparison of coil geometries under identical excitation conditions rather than device-specific predictions. The dielectric values were sourced from the IT’IS Foundation database (version 4.1) at 2.5 kHz (Hasgall et al., 2022).

To account for age-related differences in tissue composition, we explored age-dependent scaling of dielectric properties as part of a sensitivity analysis. This approach was selected to introduce physiologically motivated variations across multiple tissue types while remaining consistent with available literature on developmental dielectric changes. Simpler alternatives, such as applying uniform scaling or restricting adjustments to a subset of tissues (e.g., skull only), were not used as they do not capture known tissue-specific trends in composition and hydration.

The scaling relies on several key assumptions: (i) interspecies extrapolation from rat to human tissues, (ii) mapping of rat developmental stages to human age, and (iii) extrapolation from high-frequency dielectric measurements to the 2.5 kHz TMS regime. Each of these contributes to uncertainty in the absolute values, and their combined effect limits the interpretation of the results to relative trends rather than precise estimates. Detailed derivation and limitations of these scaling factors are provided in the Supplementary Material.

### 2.5. TMS Simulation Framework

As an illustrative example of the Thalia anatomical model, we evaluated six different coils: three figure-of-eight coils and three circular coils (Table 3). These coil geometries were selected to represent commercially available designs compatible with Magstim systems and fall within the range of typical clinical TMS coils D’Agostino et al. (2022); Zhong et al. (2025). The simulation utilized the LF magnetoquasi-static module in Sim4Life (ZMT Zurich MedTech, 2026), (version: 7.2, Grid: 1.5×1.5×1.5 mm^3^). The computational domain was discretized into tetrahedral volume elements generated from the STL surface representations of the anatomical model, rather than voxel-based (cubic) elements.

**Table 3:** TMS coils properties. All the coils proposed here are compatible with Magnetism200 (Magstim Company, 2026)

|  | Coils Name | Inside Diameter (mm) | Outside Diameter (mm) | Number of Turns |
| --- | --- | --- | --- | --- |
| 1 | Figure of 8 - 25mm [P/N 1165] | 18 (x2) | 42 (x2) | 14 (x2) |
| 2 | Figure of 8 - 50mm Prototype | 34 (x2) | 74 (x2) | 11 (x2) |
| 3 | Figure of 8 - 70mm [P/N 3190/1] | 56 (x2) | 87 (x2) | 9 (x2) |
| 4 | Circular - 50mm [P/N 9999] | 28 | 77 | 18 |
| 5 | Circular - 70mm [P/N 9762] | 40 | 94 | 15 |
| 6 | Circular - 90mm [P/N 3192/3] | 66 | 123 | 14 |

At the low frequencies used in TMS, electromagnetic interactions in biological tissue are well described by the magnetoquasi-static approximation, in which time-varying magnetic fields induce electric fields, and conduction currents dominate over displacement currents. Under this formulation, the induced electric field **E** is governed by Faraday’s law and Ohm’s law, with tissue conductivity σ as the dominant parameter, while the contribution of permittivity ε is negligible at these frequencies. The resulting linear system was solved to obtain the spatial distribution of the induced electric field within the head using Sim4Life (see Supplementary Material).

To improve the accuracy of our TMS simulations, we employed FreeSurfer to perform detailed parcellation of the gray matter (Figure S2). The coil was aligned along a line extending through the centroid of the skull and the centroid of the motor cortex parcellation and positioned approximately 7mm from the skin surface to account for the thickness of the surrounding plastic casing. For the figure-of-eight TMS coils, the alignment target was the cross-section where the two coils intersected, as this point was expected to induce the highest electric field magnitude. For circular coils, three alignment strategies were investigated: using the center of the coil, the mid-radius, and the outer edge of the coil (Figure 7 and S4). The smallest figure-of-eight coil was used as the reference configuration, and the distance between its reference point (Point D) and the corresponding points for the remaining coils was calculated to standardize anatomical positioning.

**Figure 7:**
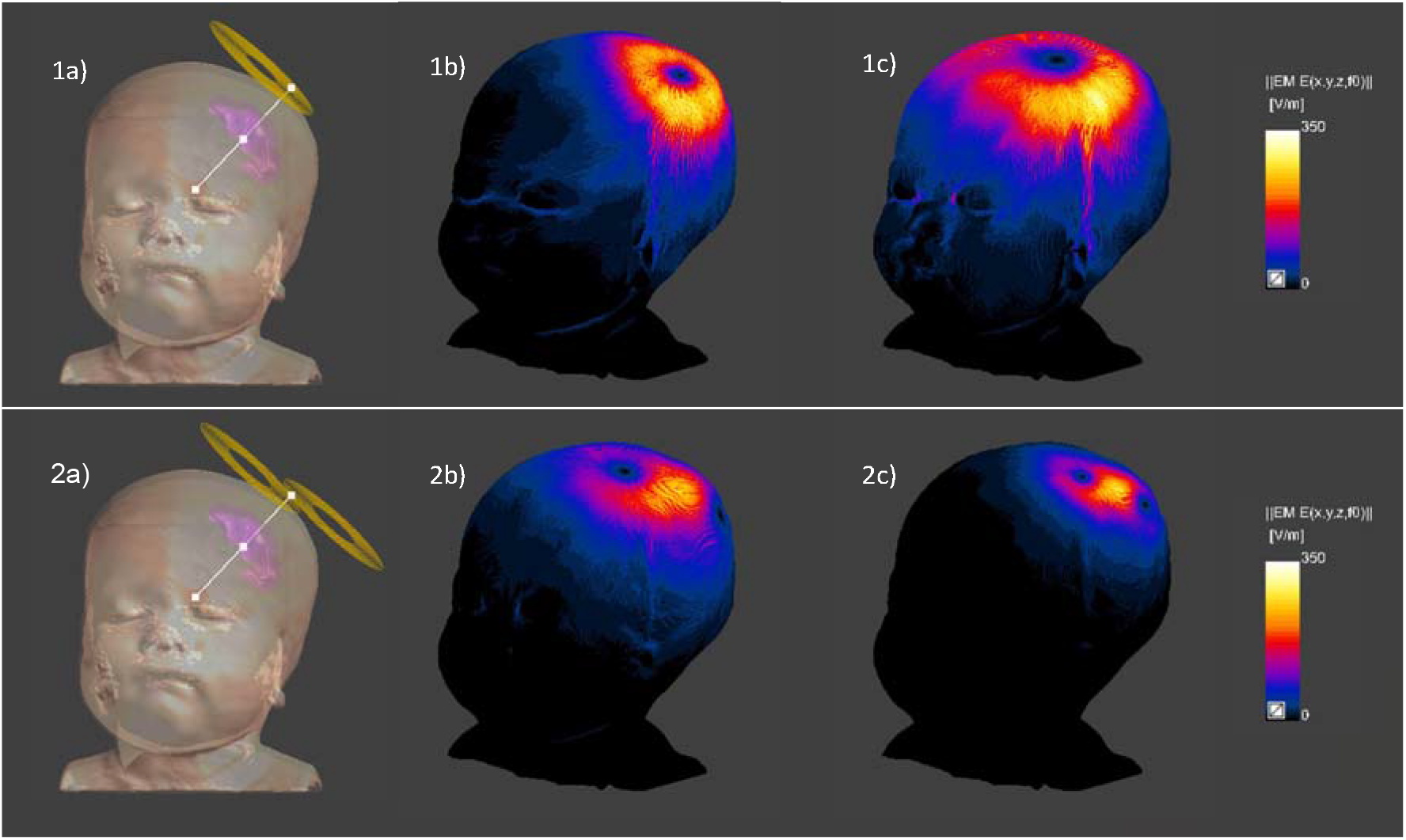
Comparison of Circular and Figure-of-Eight Coils for Illustrative Brain Gray Matter Stimulation. Top Row - Circular Coil. (1a) The mid-radius of the circular coil was aligned along a line extending through the centroid of the skull and the centroid of the motor cortex parcellation. (1b) Electric field magnitude distribution for the 50mm diameter circular coil. (1c) Electric field magnitude distribution for the 90mm diameter circular coil. **Bottom Row - Figure- of-Eight Coil** (2a) The intersection of the two windings of the figure-of-eight coil was aligned along a line extending through the centroid of the skull and the centroid of the motor cortex parcellation. (2b) Electric field magnitude distribution for the 50mm diameter figure-of-eight coil. (2c) Electric field magnitude distribution for the 25mm diameter figure-of-eight coil. Note: The same scaling (min/max values) was applied to the electric field magnitudes in subplots (1b), (1c), (2b), and (2c) to facilitate visual comparison of the simulated electric field distributions across coil configurations.

For each TMS coil, we conducted two types of simulations: one using standard dielectric properties and one incorporating dielectric properties adjusted for a 10-month-old model (sensitivity analysis). The electric field results were extracted and analyzed in MATLAB after spatial alignment with the anatomical model. We computed the maximum electric field magnitude (||Emax||) and defined a threshold equal to half of this maximum (||Emax||/2) to evaluate the volume of stimulated gray matter (V1/2) and the depth of stimulation (d1/2). A complementary analysis was performed using the spread metric (S1/2) to quantify the spatial extent of the electric field relative to its depth, following the methodology proposed by Deng et al. (2020).

## 3. Results

### 3.1. Segmentation

The segmentation process produced a comprehensive anatomical model encompassing a total of 442 distinct tissue labels, summarized in Table 4, with the complete list of segmented tissues and their corresponding volumes provided in Supplementary Table S1. Figures 4, 5, and 6 illustrate the detailed segmentation results, highlighting the accuracy achieved through both automatic and manual methods. The automated segmentation provided by FreeSurfer, complemented by manual refinements, effectively captured key brain structures and other critical anatomical features.

**Table 4:** Summary list of segmented tissues. L&R: indicates that tissues were segmented as different labels for the right and left side; C&BM: Cortex and Bone marrow segmented separately as individual labels; CNS: Central Nervous System.

|  |  |  |  |  |
| --- | --- | --- | --- | --- |
| **A. Body tissues (torso)** | Ovary (L&R) | Ankle bones (L&R) (C&BM) | Rib (L&R) (C&BM) (x12) | Cerebellar white matter (L&R) |
| Adrenal gland (L&R) | Pancreas | C1 (C&BM) | Sacrum (C&BM) | Cerebral Grey matter (L&R) |
| Air head and neck | Retina (eyes) (L&R) | C2 (C&BM) | S1-S5 (C&BM) | Cerebral White matter (L&R) |
| Arteries head & neck | Salivary glands | C3 (C&BM) | Scapulae (L&R) (C&BM) | Colliculi |
| Air abdomen (Small & Large Bowel) | Sclera (L&R) | C4 (C&BM) | Skull (C&BM) | Choroid plexuses |
| Blood Vessels Body | Skin | C5 (C&BM) | Sternum (C&BM) | Cranial Nerves other (L&R) |
| Choroid (eye) (L&R) | Small bowel contents | C6 (C&BM) | T1 (C&BM) | CSF brain |
| Ciliary muscles (eye) (L&R) | Small bowel wall | C7 (C&BM) | T2 (C&BM) | CSF spine |
| Connective tissue | Spinal Cord | Calcaneus (L&R) (C&BM) | T3 (C&BM) | Globus Pallidus (L&R) |
| Cornea (eye) (L&R) | Spleen | Carpal bones (L&R) (C&BM) | T4 (C&BM) | Hippocampus (L&R) |
| Esophagus | Stomach wall | Cartilage | T5 (C&BM) | Hypothalamus (L&R) |
| Extraocular muscles | Stomach contents | Clavicle (L&R) (C&BM) | T6 (C&BM) | Lateral Ventricle (L&R) |
| Fallopian Tubes (L&R) | Subcutaneous fat | Coccyx (C&BM) | T7 (C&BM) | Mammillary bodies |
| Gallbladder (wall & contents) | Teeth | Femur (L&R) (C&BM) | T8 (C&BM) | Medulla |
| Heart Muscle | Talus | Fibula (L&R) (C&BM) | T9 (C&BM) | Meninges Brain & Spine |
| Intrabdominal fat | Thymus | Humerus (L&R) (C&BM) | T10 (C&BM) | Midbrain |
| Intraorbital fat | Thyroid | Intervertebral disks | T11 (C&BM) | optic chiasm |
| Kidney (L & R) | Tongue | Iliac Bone (L&R) (C&BM) | T12 (C&BM) | optic nerve (L&R) |
| Large bowel contents | Trachea | Kneecap (L&R) (C&BM) | Tarsal bones (L&R) (C&BM) (x5) | Optic tract (L&R) |
| Large bowel wall | Turbinates | L1 (C&BM) | Tibia (L&R) (C&BM) | Pallidum (L&R) |
| Lens (eye) (L&R) | Urinary bladder wall | L2 (C&BM) | Ulna (L&R) (C&BM) | Pons |
| Ligaments | Uterus | L3 (C&BM) | **C. CNS Tissues** | Pituitary |
| Liver | Vagina | L4 (C&BM) | 3rd Ventricle | Putamen (L&R) |
| Lung (L&R) | Vitreous body (eye) (L&R) | L5 (C&BM) | 4th Ventricle | Septum Pellucidum |
| Lymphoid tissue (head) | Veins head & neck | Lower mandible (C&BM) | Accumbens (L&R) | Thalamus (L&R) |
| Muscles | **B. Bones** | Metacarpal bones, (L&R) (C&BM) (x5) | Amygdala (L&R) | Veins |
| Nasal Cardillage | 1st phalanges (L&R) (C&BM) (x5) | Metatarsal bones (R&L) (C&BM) | Cauda Equina | Ventral Diencephalon (L&R) |
| Nasal Mucosa | 2nd phalanges, (L&R) (C&BM) (x5) | Pelvic bone (C&BM) | Caudate (L&R) | Vermis Grey Matter |
| Ossification centers | 3rd phalanges, (L&R) (C&BM) (x5) | Radius (L&R) (C&BM) | Cerebellar grey matter (L&R) | Vermis White matter |

The brain was segmented into 36 parenchymal structures, including cortical gray matter, white matter, cerebellar components, deep gray nuclei, and brainstem structures. In addition, the meninges, the choroid plexus, and five cerebrospinal fluid (CSF) compartments were delineated, comprising the bilateral lateral ventricles, third ventricle, fourth ventricle, and remaining intracranial CSF, allowing separate quantification of total CSF and functional brain parenchymal volumes (Figure 5a-d).

The eyes and optic system were segmented into 21 distinct structures, covering ocular tissues (e.g., sclera, lens, retina, vitreous body), extraocular muscles, and optic pathways. The head and neck region contributed 15 additional segments, including air cavities, mucosa, salivary glands, tongue, skull, mandible, and associated soft tissues.

The skeletal system was represented with great anatomical detail. The upper limbs comprised 10 bones per limb, resulting in 20 segments, while the hands included 40 bones (80 segments). The lower limbs were segmented into 14 bones (28 segments), and the feet into 42 bones (84 segments) (Figure 4b). For all skeletal elements, compact bone and bone marrow were segmented separately, reflecting their distinct physical and biological properties.

The spine was segmented into 67 structures, including vertebral bodies, bone marrow, intervertebral discs, ligaments, spinal cord, and surrounding tissues (Figure 6B). The thoracic cage was divided into 50 segments, accounting for ribs and sternum, with separate compact bone and marrow compartments. Finally, 29 visceral tissues were segmented, encompassing major thoracic and abdominal organs such as the lungs, liver, kidneys, heart, spleen, gastrointestinal tract, and associated contents (Figure 6A). Together, this segmentation strategy enabled a detailed and internally consistent volumetric representation of the entire human body.

The final segmented model, exported in STL, is compatible with Sim4Life, ensuring its suitability for advanced electromagnetic simulations and further analysis.

### 3.2. Validation

To validate the model, each segmented structure was analyzed in its extracted STL format. The surface of each segment was inspected for potential isolated regions (“islands”), and manual adjustments were made as necessary to ensure precision. Additionally, all segments were independently reviewed and verified by two neuroradiologists. Feedback from the neuroradiologists was provided iteratively throughout the segmentation process, with direct comments on anatomical boundaries and tissue delineation. This led to targeted corrections across multiple structures, including refinement of gray matter boundaries and brainstem regions (Figure 3), as well as adjustments to other tissues such as extraocular muscles (Figure S1). These revisions were incorporated into the final segmentation, ensuring anatomical consistency and improving overall model accuracy.

Segmentation accuracy and reliability were further assessed by comparing morphometric measures (e.g., volume, length, or weight) with values reported in the literature. The total CSF volume within the head was calculated by summing all CSF volumes in the brain parenchyma (CSF Brain) and the volumes of the lateral (left and right), third, and fourth ventricles, resulting in 93.57 cm3. For comparison, the age-specific reference model proposed by Matsuzawa (Matsuzawa et al., 2001) provides a method to estimate theoretical CSF volume. For a 10-month-old child (0.833 years), this model yielded a brain weight of 93.89 cm3 (Eq. 1-2). The CSF volume derived from our model closely corresponds with the theoretical estimate obtained using Matsuzawa’s equations, demonstrating strong concordance between measured and reference values.

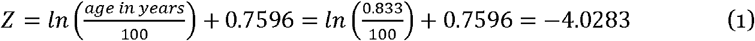

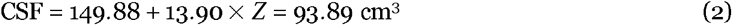

Regarding the brain parenchyma, we determined a total brain volume of 859.81cm3. Assuming a brain mass density of 1046kg/m3 (IT’IS Foundation, 2026a), this corresponds to an approximate mass of 899.36g. Based on interpolated data from Valentin et al. (2002) (Valentin, 2002), the estimated brain weight for a 10-month-old infant is 854.81 g. Furthermore, analyses of brain weight relative to age indicate that a typical 10-year-old has a brain mass ranging from 800g to 900g (Matsuzawa et al., 2021), suggesting that our estimate is consistent with values reported in the literature.

Beyond brain and CSF volumes, further validation of the pediatric model was performed by comparing measured tissue weights, organ volumes, and skeletal ratios with corresponding literature values (Table 4 5). Across all assessed organs, including the lungs, heart, liver, kidneys, pancreas, spleen, stomach, thymus, and ovaries, the measured weights and volumes fell within the ranges reported in previous studies (Chang et al., 2021; Kelsey et al., 2021; Konuş et al., 2013). Similarly, skeletal metrics, such as bone lengths and inter-bone ratios (e.g., radius/humerus, tibia/femur, humerus/femur, radius/tibia), closely matched normative values (Robinow and Chumlea, 1982; Weaver et al., 2014), confirming the anatomical plausibility of the reconstructed model. The overall agreement between measured and reference values across multiple organ systems and skeletal proportions provides additional confidence in the accuracy and reliability of the segmented pediatric model.

**Table 5:** Validation of the segmented pediatric model. Quantitative comparison between measured tissue weights, volumes, and bone length ratios from the segmented pediatric model and corresponding literature reference values. Agreement across tissues and skeletal ratios confirms the validity of the reconstructed model.

| Tissue | Measurement type | Measured value | Literature value |
| --- | --- | --- | --- |
| Lung (R) | Weight | 112.5 g | 70–180 g (Chang et al., 2021) |
| Lung (L) | Weight | 102.3 g | 70–180 g (Chang et al., 2021) |
| Brain | Weight | 899.4 g | 800–900 g (Chang et al., 2021) Matsuzawa et al. (2021) |
| Brain CSF | Volume | 93.6 cm<sup>3</sup> | 93.8 cm<sup>3</sup> (Matsuzawa et al., 2001) |
| Heart | Weight | 48.3 g | 25–50 g (Chang et al., 2021) |
| Kidneys | Weight | 48.4 g | 45–80 g (Chang et al., 2021) |
| Kidney (R) | Length (longitudinal) | 56.2 mm | 52–66 mm (Konu, s et al., 2013) |
| Kidney (L) | Length (longitudinal) | 56.4 mm | 54–68 mm (Konu, s et al., 2013) |
| Liver | Weight | 257.6 g | 220–380 g (Chang et al., 2021) |
| Liver | Length (longitudinal) | 74.4 mm | 70–90 mm (Konu, s et al., 2013) |
| Pancreas | Weight | 8.5 g | 7–17 g (Chang et al., 2021) |
| Spleen | Weight | 30.3 g | 15–32 g (Chang et al., 2021) |
| Spleen | Length (longitudinal) | 67 mm | 53–74 mm (Konu, s et al., 2013) |
| Stomach | Weight | 14.2 g | 12–23 g (Chang et al., 2021) |
| Thymus | Weight | 22.1 g | 21–34 g (Chang et al., 2021) |
| Ovaries | Volume | 0.51 cm<sup>3</sup> | 0.1–1.5 cm<sup>3</sup> (Kelsey et al., 2021) |
| Sternal bone | Length | 6.7 cm | 5.7 cm (Weaver et al., 2014) |
| Radius/<br>Humerus | Bone length ratio | 76.6 | 0.73–0.85<br>(Robinow and Chumlea, 1982) |
| Tibia/<br>Femur | Bone length ratio | 82 | 0.75–0.87<br>(Robinow and Chumlea, 1982) |
| Humerus/<br>Femur | Bone length ratio | 77.3 | 0.73–0.85<br>(Robinow and Chumlea, 1982) |
| Radius/<br>Tibia | Bone length ratio | 72.2 | 0.71–0.83<br>(Robinow and Chumlea, 1982) |

### 3.3. Tissue Properties

The tissue properties used in the standard simulations were based on the IT’IS database (IT’IS Foundation, 2026a), which primarily reflects adult tissue characteristics. However, since our study focuses on a pediatric subject, it was necessary to adjust these properties to be age appropriate. To investigate the potential impact of such differences, we conducted a sensitivity analysis using age-dependent scaling ratios rather than attempting to define exact pediatric dielectric properties. To achieve this, we determined age-dependent ratios using methodologies from Martin’s study (Jeong et al., 2021) and Athena’s model (Ntolkeras et al., 2023). Data from Peyman et al.’s study (with corrections) (Peyman et al., 2002) were extrapolated to obtain permittivity and conductivity values for a 2.75-day-old rat (Supplementary Figure S3). Ratios of permittivity and conductivity were then calculated by comparing values at 2.75 days to those at 70 days, analogous to the 10-month-old/adult human ratio.

Among the tissues analyzed, the skull (cortical bone) exhibited the highest age-dependent scaling ratios, with a relative permittivity ratio of 2.04 and an electrical conductivity ratio of 3.50. For the skin and brain, the conversion ratios were 1.67 and 1.43 for permittivity, and 1.94 and 1.65 for electrical conductivity, respectively. We also calculated the ratios for muscle, gland, and liver tissues. To estimate the scaling factor for soft tissues not explicitly reported, we used an average scaling ratio derived from four soft tissues, excluding the skin and skull (cortical bone), which were treated as outliers (Jeong et al., 2021). This yielded an average permittivity ratio of 1.32 and a conductivity ratio of 1.55. These ratios were utilized to obtain the dielectric properties (Table S2) needed for the subsequent sensitivity analysis. Consistent with the assumptions outlined above, these values are interpreted as representative variations to assess model sensitivity rather than definitive pediatric tissue properties.

### 3.4. Illustrative Example: TMS Simulation Results

Table 6 summarizes the distances between the reference point (Point D) of the smallest figure-of-eight coil and the corresponding reference points obtained using the three alignment strategies for the circular coils. Among the three alignment strategies evaluated, the mid-radius alignment resulted in the shortest distances and was therefore used for the subsequent circular-coil simulations. Table 7 summarizes the peak electric field (||Emax||), stimulated gray matter volume (V1/2), stimulation depth (d1/2), and spread (S1/2) for the six coil configurations. Corresponding visualizations are provided in the Supplementary Material.

**Table 6:** Distances Used to Compare Circular Coil Alignment Strategies. This table presents the distances from each coil’s point D (the deepest point where the electric field exceeds Emax/2) to the reference point D on Coil 1. Distances are shown for simulations using standard dielectric properties, adult dielectric properties, and age-adjusted dielectric properties. For each circular coil, the smallest distance is highlighted in bold and was used for the subsequent simulations.

|  | Coil Type | Coil Size | Coil Position | Distance [mm]<br>Standard | Distance [mm]<br>Age Adjusted |
| --- | --- | --- | --- | --- | --- |
| 1 | Figure-of-8 | 25mm x 2 | Cross | 0 (Ref) | 0 (Ref) |
| 2 | Figure-of-8 | 50mm x 2 | Cross | 13.89 | 1.53 |
| 3 | Figure-of-8 | 70mm x 2 | Cross | 6.17 | 6.54 |
| 4 | Circular | 50mm | Center | 38.80 | 39.32 |
|  |  |  | Mid-Radius | **19.23** | **19.23** |
|  |  |  | Outer | 42.22 | 38.24 |
| 5 | Circular | 70mm | Center | 52.30 | 41.62 |
|  |  |  | Mid-Radius | **5.89** | **1.82** |
|  |  |  | Outer | 9.50 | 9.50 |
| 6 | Circular | 90mm | Center | 42.86 | 31.40 |
|  |  |  | Mid-Radius | **17.19** | **5.48** |
|  |  |  | Outer | 51.02 | 51.02 |

**Table 7:** Summary of electric field metrics for the illustrative TMS simulations. This table summarizes the electric field metrics obtained from the illustrative transcranial magnetic stimulation (TMS) simulations presented in Figure 8. The columns include the coil name, maximum electric field strength (Emax), the volume (V1/2) above the Emax/2 threshold, the half-value depth (d1/2), and the half-value spread (S1/2). Results are reported for figure-of-eight and circular coils of different sizes using both standard dielectric properties and age-adjusted dielectric properties (sensitivity analysis).

| | Coil Name | $E_{max}$ | $V_{1/2}$ | $d_{1/2}$ | $S_{1/2}$ |
| --- | --- | --- | --- | --- | --- |
| Standard | 1 figure-of-eight coil (25mm) | 374.07 | 11.55 | 1.12 | 10.35 |
|  | 2 figure-of-eight coil (50mm) | 727.02 | 2.59 | 1.11 | 2.34 |
|  | 3 figure-of-eight coil (70mm) | 439.28 | 94.44 | 1.88 | 50.10 |
|  | 4 circular coil (50mm) | 671.72 | 11.29 | 1.00 | 11.28 |
|  | 5 circular coil (70mm) | 509.76 | 90.42 | 1.53 | 59.06 |
|  | 6 circular coil (90mm) | 600.29 | 92.11 | 1.50 | 61.39 |
| Age-Adjusted | 1 figure-of-eight coil (25mm) | 424.40 | 9.47 | 1.12 | 8.49 |
|  | 2 figure-of-eight coil (50mm) | 946.62 | 1.56 | 0.99 | 1.57 |
|  | 3 figure-of-eight coil (70mm) | 498.72 | 77.06 | 1.65 | 46.65 |
|  | 4 circular coil (50mm) | 881.66 | 5.58 | 1.00 | 5.58 |
|  | 5 circular coil (70mm) | 606.53 | 54.87 | 1.50 | 36.57 |
|  | 6 circular coil (90mm) | 734.22 | 48.39 | 1.23 | 39.25 |

The simulated peak electric field magnitude (||Emax||) exceeded 300 V/m for all coil configurations under both standard and age-adjusted dielectric properties (Figure 8). Moreover, Figure 8 shows that a small 25mm figure-of-eight coil produces highly focal stimulation with comparatively lower peak electric field values, whereas the 90 mm circular coil provides generated a higher Emax together with a broader stimulated gray matter volume. The 70 mm circular and figure-of-eight exhibited intermediate characteristics. Among the simulated configurations, the 50 mm figure-of-eight coil combined relatively high peak electric field values with a comparatively small stimulated gray matter volume.

**Figure 8:**
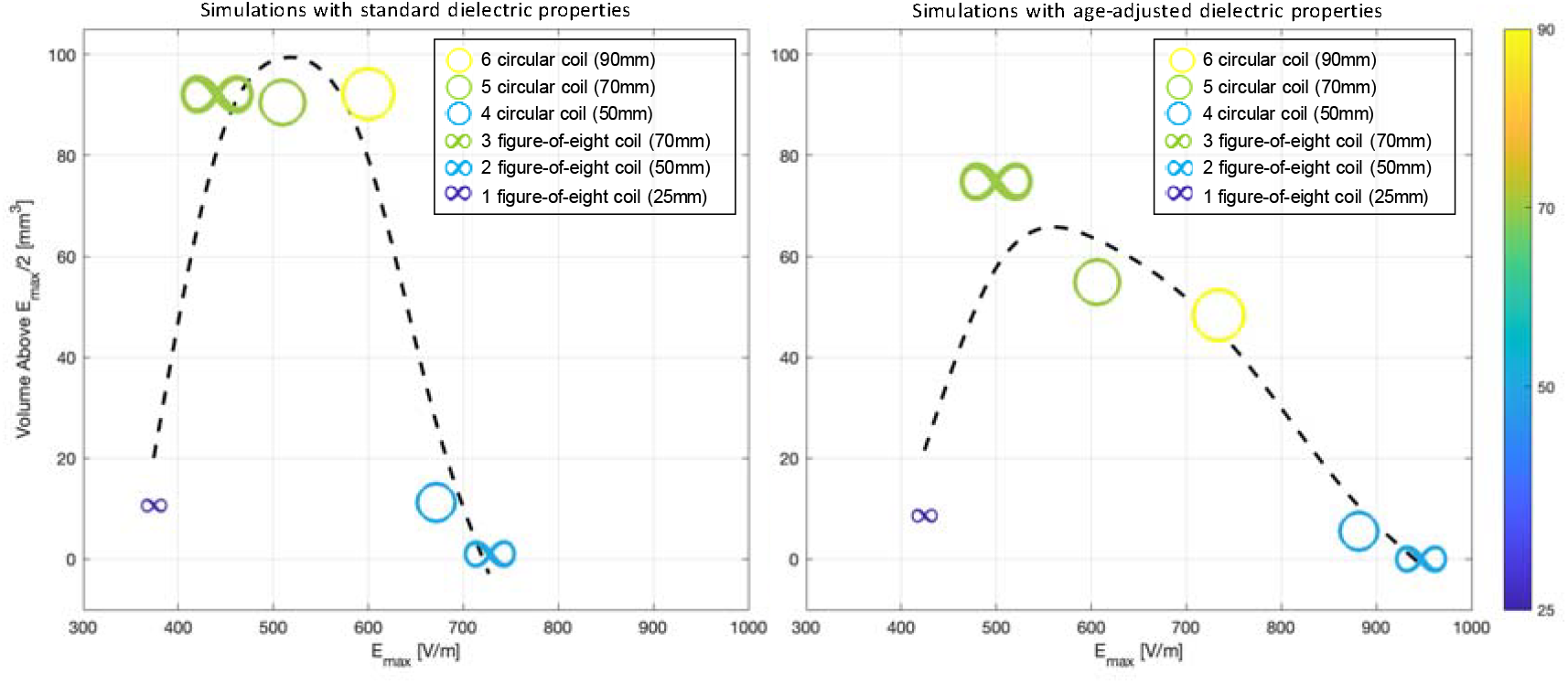
Electric Field Max vs. Volume Above Threshold for TMS Coils. Comparison of simulation results for different types of TMS coils. The two subplots show the relationship between the maximum electric field magnitude *E*_*max*_ in V/m (x-axis) and the volume above the *E*_*max*_*/*2 threshold in mm^2^ (y-axis). On the left, results are presented with standard dielectric properties; on the right, with age-adjusted dielectric properties. Both plots share the same axes and limits. The six coil types compared are: (i) three circular coils of different sizes, represented by circles of varying sizes and colors corresponding to the coil size; (ii) three figure-of-eight coils, represented by infinity symbols of different sizes and colors according to the coil size. A smoothing spline is plotted to illustrate the general trend for the six coils in each subplot.

Simulations using age-adjusted dielectric properties showed similar trends across the evaluated coil configurations. Higher peak electric field magnitudes were observed compared with the standard dielectric-property simulations. This difference is consistent with the increased conductivity, particularly in the skull, assumed in the sensitivity analysis, which reduces attenuation of induced currents and results in higher simulated electric field magnitudes. Consequently, increasing the threshold (||Emax||/2) reduces the corresponding stimulated gray matter volume. The complementary analyses of stimulation depth and spatial spread (Supplementary Material) showed trends consistent with the primary electric field analyses across the simulated coil configurations.

## 4. Discussion

### 4.1. Numerical Model Acquisition

The Thalia model is a comprehensive computational tool designed to provide a detailed anatomical representation of a full body model, primarily intended for studying tissue interactions with electromagnetic fields generated by medical devices and conducting dosimetry studies in 10-month-old children. One of the model’s significant advantages is its development based on in-vivo medical imaging of a 10-month-old child, rather than relying on morphing from another model of a different age or gender, as seen with models like Nina of the GSF Voxel Family (Petoussi-Henss et al., 2002), which was morphed from Roberta of the same family. Importantly, all scans were obtained from the same child. In contrast, other models, such as the UF Family (Lee et al., 2010) and the XCAT (Segars et al., 2015), used scans with limited body coverage and filled the gaps with data from different subjects, or based their segmentation on CT scans.

Electric fields generated by medical devices and conducting dosimetry studies in 10-month-old children. One of the model’s significant advantages is its development based on in-vivo medical imaging of a 10-month-old child, rather than relying on morphing from another model of a different age or gender, as seen with models like Nina of the GSF Voxel Family (Petoussi-Henss et al., 2002), which was morphed from Roberta of the same family. Importantly, all scans were obtained from the same child. In contrast, other models, such as the UF Family (Lee et al., 2010) and the XCAT (Segars et al., 2015), used scans with limited body coverage and filled the gaps with data from different subjects or based their segmentation on CT scans.

The use of multiple MRI sequences enabled precise segmentation across tissues, particularly in the head and neck regions where anatomical detail is critical for neuroimaging and bioelectromagnetic applications. With a total of 442 segmented tissues, Thalia represents a high-resolution model compared to existing pediatric phantoms, such as MARTIN (114 tissues) and ATHENA (267 tissues), while maintaining full-body anatomical consistency. It also enabled precise segmentation. For instance, T1 MPRAGE served as the basis for automated brain segmentation, while T2 FLAIR allowed for refined delineation of fluids, such as veins, arteries, and CSF. The inclusion of a sagittal IR sequence of the spine was paramount in achieving precise vertebral segmentation. Additional scans, such as the IR and T1 full-body sequences, were used to segment other body parts, including extremities (e.g., limbs). More specialized scans, like the T2 HASTE and IR sequences of the chest, assisted in the accurate segmentation of organs (e.g., lungs, heart). With a total of 442 segments, the Thalia model can be considered high-resolution. For comparison, the MARTIN model (Jeong et al., 2021), developed in 2021 and added to the virtual population (IT’IS Foundation, 2026a), consisted of 28 head and 86 body tissues. The Thalia model aims to achieve a high level of resolution, similar to the ATHENA model from 2022 (Ntolkeras et al., 2023), which includes a total of 267 tissues.

This level of anatomical detail enables more accurate assignment of tissue-specific properties and provides a flexible platform for a wide range of computational studies, including electromagnetic simulations, dosimetry, and device design in pediatric populations.

#### 4.1.1. Image Acquisition, Preprocessing, and Registration

The resolution of the MRI scans was not always optimal for detailed analysis, posing significant challenges despite our efforts to standardize all images to a resolution of 0.5 × 0.5 × 0.5 kg/mm3. This resampling process, while intended to enhance uniformity and comparability across datasets, could not entirely overcome the inherent limitations of the original scans. For example, the full-body scans had relatively low in-plane resolutions of 1.25 × 1.25 kg/mm2 and 0.73 × 0.73 kg/mm2, coupled with thick slices of 5.0 mm. When these images were resampled, the increased voxel count did not equate to greater anatomical detail but instead introduced potential interpolation artifacts and image blurring. This issue was particularly pronounced in the spine and neck and chest scans, where the original slice thicknesses of 3.0 mm and 5.0 mm significantly constrained the achievable level of detail. Although resampling improved resolution along the slice direction, the original limitations persisted, thereby diminishing thepotential gains from this enhancement. In contrast, the brain scans, particularly the T2 FLAIR sequence, originally had a finer in-plane resolution of 0.39 × 0.39 kg/mm2, benefiting more from resampling, especially in improving through-plane resolution. However, even in these cases, the original anisotropy where the resolution in one direction was markedly different from the others, meant that the final images still retained some degree of distortion and loss of detail.

Using multiple MRI scans for segmentation provided a comprehensive view of the anatomy, yet it introduced significant challenges, particularly in image co-registration. Achieving precise alignment of images from different sessions was crucial, as even minor discrepancies could lead to shifts that compromised segmentation accuracy. This challenge was exacerbated by the diversity of MRI scans, each targeting different body regions (such as the brain, chest, and spine) with distinct anatomical landmarks (Figure 1). Furthermore, the use of varying imaging modalities and parameters for each body part added another layer of complexity. Differences in resolution, field-of-view, and slice thickness demanded sophisticated image processing techniques to effectively harmonize the datasets. Despite these obstacles, the coregistration process was successfully completed, allowing us to create an accurate three-dimensional (3D) model of the infant’s anatomy.

#### 4.1.2. Segmentation and Validation

Regarding the brain tissues, the segmentation process employed both automated methods using the infant-specific FreeSurfer framework (Zöllei et al., 2020), as well as the semi-automated and manual refinement to achieve high-quality results. FreeSurfer’s automated segmentation provided a solid foundation by accurately delineating anatomical structures through sophisticated algorithms. However, semi-automated and manual refinements were essential for correcting minor inaccuracies and further enhancing the overall detail and precision of the segmentation by refining the initial output and adding new tissues (e.g., pituitary, hypothalamus, colliculi, mammillary bodies, ventricles, CSF, vessels, etc.). The choice of MRI sequences played a crucial role in this process, with each contributing to different aspects of image quality and detail. For brain segmentation, high-resolution scans like the AX T1 MPRAGE were particularly effective. This sequence, with a voxel size of 0.94 × 0.94 × 1.0 mm kg/mm3 and a short inversion time (TI) of 958 ms, which allowed for clear visualization of gray and white matter while minimizing CSF signal interference. The high resolution and specific TI setting were instrumental in accurately delineating brain regions, ensuring precise measurements of brain volume. In contrast, segmentation of CSF benefited from the AX T2 FLAIR FS sequence. This scan, with a voxel size of 0.39 × 0.39 × 3.0 mm kg/mm3 and a longer TI of 2500 ms, was optimized to suppress the CSF signal, enhancing the contrast between CSF and surrounding brain tissues. Overall, these methods allowed us to successfully segment brain tissues from CSF and compute brain weight and CSF volume.

For the segmentation of other head tissues, the ability to use three orthogonal views (axial, coronal, and sagittal) for each MRI sequence was invaluable. This approach enabled us to consistently select the best view for delineating tissue boundaries during the segmentation process. Additionally, the 3D visualization tools provided by 3D Slicer were instrumental in validating the segmentation by allowing us to compare the tissue shapes with their expected anatomical structures. This was particularly useful for tissues with distinctive shapes, such as the skull, lower mandible, and facial skin. The use of 3D visualization in these cases significantly aided in verifying that the segmentation progressed accurately.

#### 4.1.3. Limitations

First, the resolution of the MRI scans was not always optimal, making precise anatomical segmentation challenging. Using MRI scans from different body parts also complicated accurate image co-registration. Additionally, our segmentation process relied exclusively on MRI data. While MRI is preferred for imaging very young children due to its non-invasive nature and absence of ionizing radiation, it provides lower spatial resolution compared to other techniques, such as CT. To address them, we iteratively refined the segmentations using predefined validation metrics, continuously reviewing and adjusting the images to improve accuracy and ensure proper anatomical alignment.

Furthermore, although segmentation accuracy was carefully validated through expert review and comparison with literature values, small anatomical inaccuracies may still persist, particularly in regions with limited image resolution or contrast. These limitations should be considered when interpreting simulation results derived from the model.

### 4.2. Illustrative Example: TM Simulation Discussion

A qualitative comparison with published adult TMS modeling studies provides important context for the illustrative application presented here. Zhong et al. (2025) analyzed induced electric fields across 50 heterogeneous adult head models (ages 22–35) using finite-element simulations in Sim4Life under comparable excitation conditions (5 kA, 2.5 kHz) and similar commercial coil configurations. Their analysis, based on Emax and V1/2 metrics as well, shows that figure-of-eight coils provide higher focality, while circular coils generate broader stimulation patterns, consistent with the trends observed in the present simulations. They also report substantial intersubject variability, often exceeding differences between coil designs. However, the electric field magnitudes reported in adult models (typically ∼100– 300 V/m) are substantially lower than those observed in our pediatric simulations. In our model, Emax values range from ∼300–700 V/m under standard conditions and ∼400–900 V/m for the sensitivity analysis which incorporates age-adjusted dielectric properties. This comparison remains indirect because the simulations were performed using different anatomical models and computational assumptions. Consequently, the observed differences in electric field magnitude should not be attributed solely to developmental anatomy but instead provide contextual comparison with previously published studies.

The TMS simulations presented in this work should be interpreted an illustrative example of the Thalia anatomical model, demonstrating one potential use of this computational resource rather than providing definitive guidance for pediatric TMS. All coils were evaluated under identical excitation conditions, and coil placement was standardized to ensure consistent geometric alignment, rather than to replicate neuronavigated clinical positioning. Consequently, the results should be interpreted as relative comparisons within the simulated framework and not as evidence for optimal coil selection or clinical recommendations.

Finally, age-dependent dielectric properties were incorporated as part of a sensitivity analysis based on literature-derived scaling, rather than direct pediatric measurements. While the exact scaling factors remain uncertain, the overall direction of variation is supported by known developmental differences in tissue composition, including higher hydration and conductivity in younger populations. Accordingly, these variations should be interpreted as plausible bounds for assessing model sensitivity rather than direct representations of infant tissue properties. Importantly, the reported electric field values represent biophysical metrics and should not be interpreted as direct indicators of clinical safety or tolerability, as these depend on additional factors such as waveform characteristics, individual anatomical variability, and subject-specific neurophysiological responses. The standardized excitation, simplified coil positioning, and absence of experimental or clinical validation further limit the interpretation of the simulated electric fields.

## 5. Conclusion

This study presents Thalia, a validated, high-resolution computational model of a 10-month-old female comprising 442 distinct tissues, addressing a critical gap in pediatric anatomical phantoms. By leveraging in vivo MRI data and extensive automatic and semi-automatic expert-guided segmentation, the model achieves a level of anatomical fidelity that supports detailed electromagnetic simulations, particularly in the head and neck regions. Quantitative validation against age-matched anatomical reference data confirms the accuracy and reliability of the reconstructed anatomy. Importantly, the model is entirely non-morphed and derived from a single subject, ensuring anatomical consistency across the whole body. This approach overcomes limitations of existing pediatric models that rely on morphing or partial datasets and provides a more realistic representation of early developmental anatomy.

Beyond its anatomical detail, Thalia integrates 442 segmented tissues across the entire body, including fine brain structures, vasculature, skeletal components, and organs, enabling multi-scale and multiorgan investigations. This level of detail allows for more accurate assignment of tissue-specific properties and supports a wide range of computational studies requiring precise anatomical and biophysical realism.

Overall, this work establishes Thalia as a powerful and versatile platform for in silico investigations in pediatric populations. In addition to neuromodulation studies, the model is well suited for application such as dosimetry, safety assessment, medical device design, and the study of developmental anatomical variability. Its high resolution and validated anatomical accuracy make it particularly valuable for scenarios where small-scale anatomical differences can significantly influence simulation outcomes. While the present simulations provide critical insight into electric field distributions and coil selection in infants, future work should extend the use of this model to additional modalities and incorporate experimental and clinical validation to further bridge the gap between simulation and practice. Furthermore, expanding this framework to include additional subjects and age groups could support the development of a comprehensive library of pediatric models, enabling population-based analyses and improving the generalizability of computational studies.

## Acknowledgments

Authors acknowledge ‘Sim4Life by ZMT, www.zurichmeditech.com‘ for Science License. The research reported in this publication was supported by the United States National Institutes of Health under the National Institute of Biomedical Imaging and Bioengineering (NIBIB) award number NIH/NIBIB: R01EB024343

## Credit authorship contribution statement

**Alice Albrecht:** Writing – review & editing, Writing – original draft, Visualization, Methodology, Investigation, Formal analysis, Data curation, Software. **Georgios Ntolkeras**: Writing – review & editing, Software, Methodology, Investigation, Formal analysis, Data curation, Conceptualization. **Lilla Zöllei:** Writing – review & editing, Investigation. **Georgios A. Sideris:** Writing – review & editing, Supervision, Data curation. **Francesca Marturano:** Writing – review & editing, Data curation, Supervision. Michael H. Lev: Writing – review & editing, Supervision, Resources. **Ellen Grant:** Writing – review & editing, Resources, Methodology, Funding acquisition, Conceptualization. **Giorgio Bonmassar:** Writing – review & editing, Writing – original draft, Resources, Methodology, Funding acquisition, Conceptualization.

## Data Availability

The Thalia model will be made available after publication through the Analogue Brain Imaging Laboratory (ABILAB) at the Athinoula A. Martinos Center for Biomedical Imaging, and through the IT’IS Foundation Virtual Population platform (IT’IS Foundation, 2026b), alongside established models such as Athena and Martin.

The released data include the full anatomical segmentation derived from high-resolution MRI (0.5 × 0.5 × 0.5 mm3), associated tissue properties, and surface representations of all tissues (e.g., STL files). These data are provided in a consistent anatomical coordinate system aligned with the whole-body T1-weighted MRI reference space, ensuring reproducibility and interoperability across studies.

A simulation-ready version of the model will also be accessible through the IT’IS Virtual Population framework, enabling direct use in electromagnetic simulation environments such as Sim4Life. In this context, computational meshes (e.g., tetrahedral volume meshes for FEM simulations) are generated within the simulation environment from the provided anatomical model. The anatomical model itself is freely available. Simulation-ready versions distributed through the IT’IS platform are accessible in accordance with their standard access policies (e.g., directly available to Sim4Life users or at a minimal processing cost for non-users).

## Notes

### Competing Interest Statement

The authors have declared no competing interest.

