## Supplementary Material for "Development and Validation of Thalia: A High-Resolution Pediatric Computational Model of a 10-Month-Old Infant"

#### Feedback on the segmentation

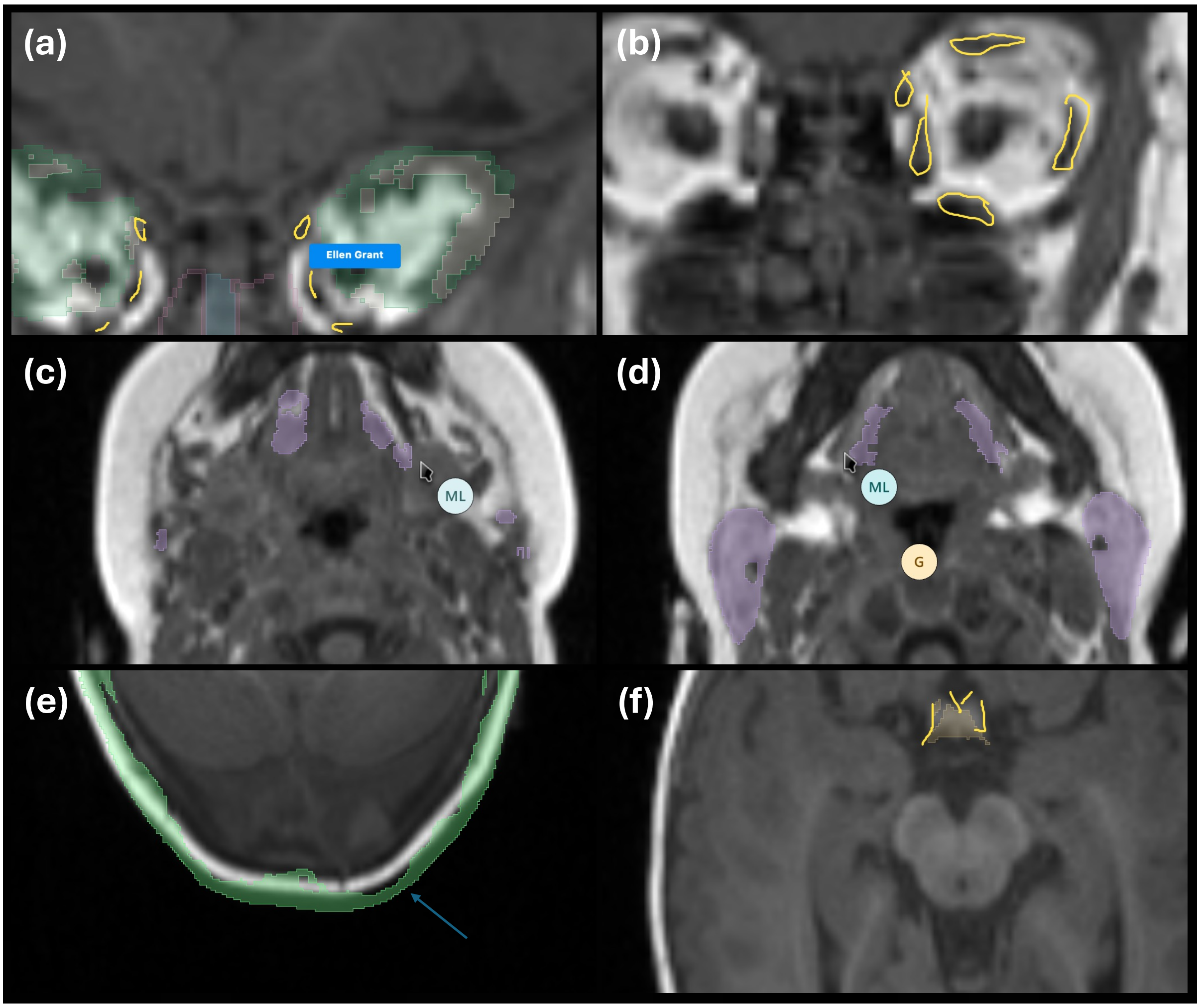


**Figure S1: Feedback example for segmentation validation.** Feedback on the segmentation was provided by two subspecialized different neuroradiologists and their recommendations were incorporated in the final version of the model. (a,b) yellow lines indicate the extraocular muscles identified by P.E.G. ("Ellen Grant" in the figure). (c,d) Feedback on the segmentation of the parotids and the sublingual salivary glands by M.H.L. ("ML" in the figure). (e) arrow pointing at the occipital regions of the skull that appeared to be outside the expected physiologic limits based on the source T1 MRI images as outlined by P.E.G and M.H.L. (f) yellow lines indicate the optic chiasm as outlined by P.E.G.

#### Parcellation of Brain Gray Matter

The brain gray matter parcellation provided by FreeSurfer (Fischl, 2012) offers a refined mapping of cortical regions within the brain (Figure S2). This high-resolution data enables precise identification and localization of specific areas, such as the precentral gyrus (motor cortex) and the postcentral gyrus (somatosensory cortex). Targeting these regions with TMS (Figure S4) allows for more accurate and targeted stimulation.

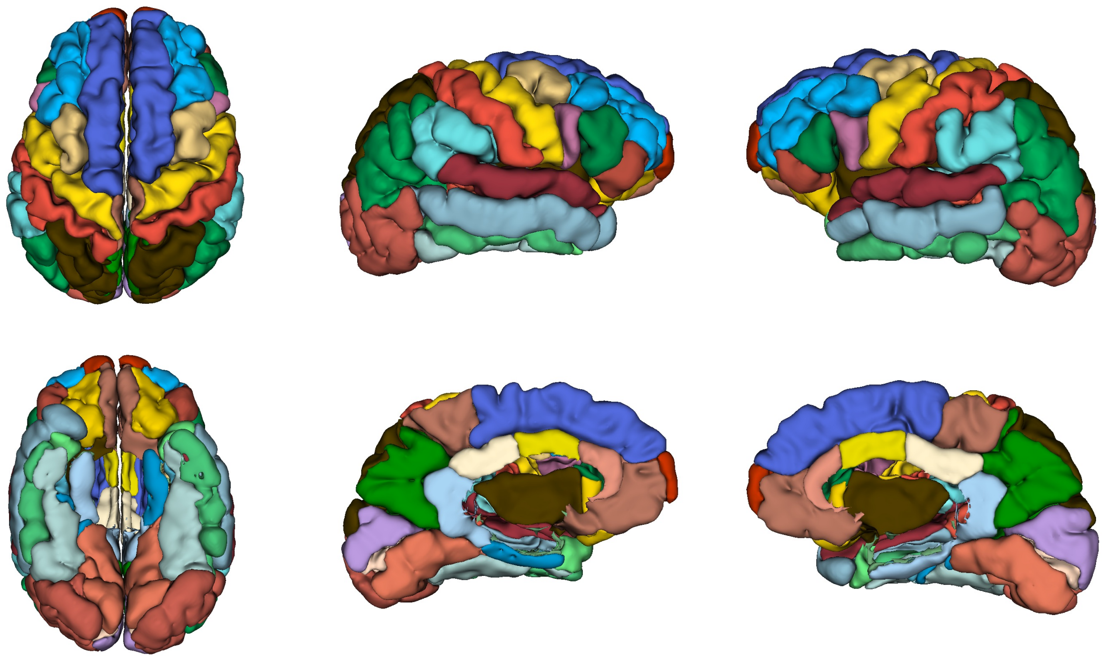

**Figure S2: High-Resolution Brain Gray Matter Parcellation by FreeSurfer.** Axial and coronal views of the gray matter parcellation provided by FreeSurfer, highlighting the cortical regions of the brain. (a) Axial view from the superior aspect (top-down), (b) Axial view from the inferior aspect (bottom-up), (c) Coronal view of the right hemisphere from the lateral aspect (outside), (d) Coronal view of the right hemisphere from the medial aspect (inside), (e) Coronal view of the left hemisphere from the lateral aspect (outside), and (f) Coronal view of the left hemisphere from the medial aspect (inside).

### Tissues Properties

Studies on various animal models, including mice (Peyman et al., 2001), rats (Gabriel, 2005), and pigs (Peyman et al., 2009), demonstrate that dielectric properties are significantly influenced by age due to changes in tissue composition and hydration levels. As individuals age, tissue fat increases while muscle mass decreases, altering both the dielectric constant and conductivity. Additionally, tissue hydration levels, which affect dielectric properties, generally decrease with age. To address these age-related changes, we implemented an exploratory scaling approach used solely for sensitivity analysis purposes.

While numerous studies exist, many focus on high frequencies, such as studies in pigs Peyman et al. (2009). Gabriel (2005) study covers a frequency range from 300 kHz to 300 MHz but provides data only for brain and skull tissues, starting from 10 days-old rat, which is roughly equivalent to 3 years in humans. *Thalia* represents a 10-month-old human, corresponding to approximately 2.75 rat days during the pre-pubertal phase (Sengupta, 2013).

The adopted scaling framework is based on three main assumptions: (i) that relative age-dependent trends observed in rat tissues are qualitatively transferable to human tissues, (ii) that developmental time can be approximated across species using established mappings, and (iii) that relative changes in dielectric properties with age are not strongly frequency-dependent, allowing extrapolation from high-frequency measurements to the 2.5 kHz regime. Each of these assumptions introduces uncertainty, with the combined effect likely leading to substantial variability in the absolute dielectric values.

Despite these limitations, we chose this approach over simpler alternatives for two main reasons. First, it preserves tissue-specific age effects (e.g., differences between skull and soft tissue), which are not captured by uniform scaling methods. Second, it is grounded in existing experimental literature. In the case of low-frequency (LF) data, the available literature is very limited, and the study by Gabriel (2005) remains one of the few comprehensive references. Therefore, we preferred to base our approach on published data rather than applying arbitrary adjustments. Approaches focusing on a single tissue (e.g., skull-only scaling) or using narrow adjustment ranges were considered less appropriate, as they may underestimate the variability in electric field distributions across tissues.

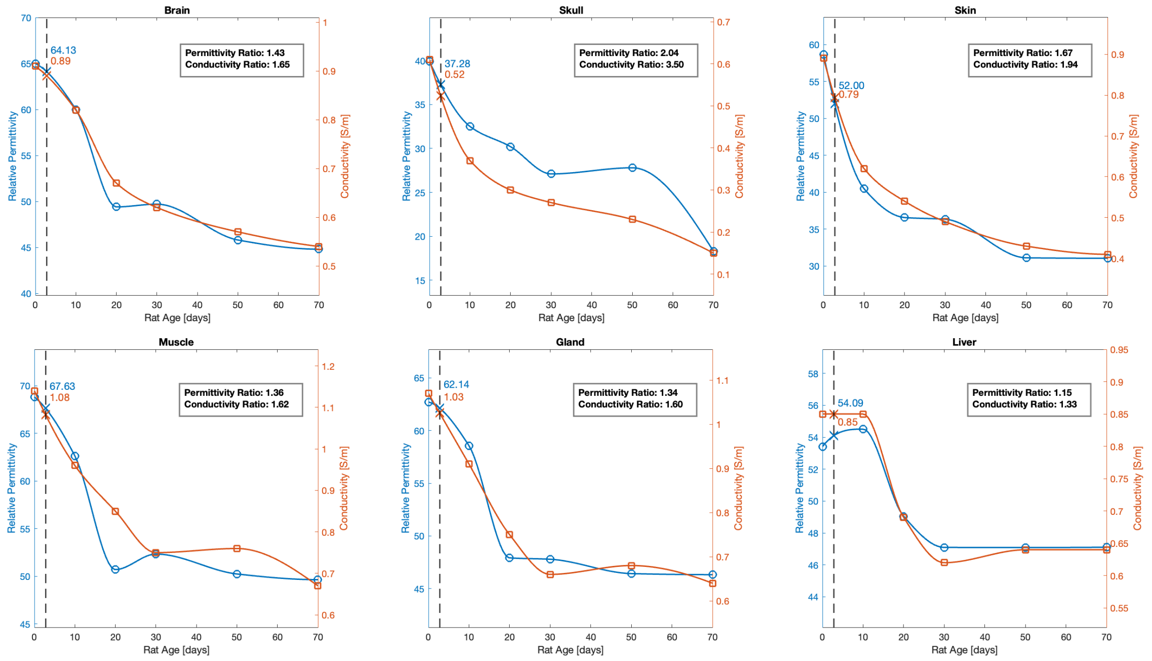


**Figure S3: Age-Dependent Dielectric Property Interpolation for Various Organs.** Six subplots display the dielectric properties of brain, skull (cortical bone), skin, muscle, gland, and liver, arranged from top left to bottom right. The X-axis shows rat age in days, with two curves plotted for each organ: relative permittivity (left Y-axis, blue) and electrical conductivity (right Y-axis, orange). Dashed line correspond to 2.75 rat days, representing the equivalent of a 10-month-old human. Ratios of permittivity and conductivity were calculated by comparing the values at 2.75 days to those at 70 days, analogous to the 10-month/adult human ratio.

### Electromagnetic Formulation and FEM Implementation

Transcranial Magnetic Stimulation (TMS) was modeled in Sim4Life using a magnetoquasi-static (MQS) formulation, in which time-varying magnetic fields generated by the coil induce electric fields in biological tissues. This approximation is appropriate for the frequency content of TMS pulses, where conduction currents dominate over displacement currents.

Under the MQS formulation, the electric field $\mathbf{E}$ is expressed as:

$$\mathbf{E} = -\frac{\partial \mathbf{A}}{\partial t} - \nabla \phi, \tag{1}$$

where $\mathbf{A}$ is the magnetic vector potential and $\phi$ is the scalar electric potential. The inductive term $-\frac{\partial \mathbf{A}}{\partial t}$ represents the primary mechanism of field generation in TMS, while the potential term accounts for charge redistribution at tissue boundaries due to conductivity differences.

The governing equation is derived from Ampère's law neglecting displacement currents:

$$\nabla \times \left( \frac{1}{\mu} \nabla \times \mathbf{A} \right) = \mathbf{J}, \tag{2}$$

where $\mathbf{J} = \sigma \mathbf{E}$ is the conduction current density. This formulation captures the inductive coupling between the TMS coil and the head tissues.

In Sim4Life, the coil is modeled as a current-driven source with a prescribed time-varying waveform. The resulting electromagnetic problem is solved using the finite element method (FEM) on a tetrahedral mesh, with tissue-specific electrical conductivity $\sigma$ and magnetic permeability $\mu$ assigned to each compartment. Boundary conditions are defined to ensure a well-posed problem, typically assuming electrically insulating outer boundaries.

Although both conductivity ($\sigma$) and permittivity ($\varepsilon$) are defined as material properties, the contribution of permittivity is negligible in the MQS regime at TMS frequencies. Therefore, displacement currents are not modeled, and the system effectively behaves as a conductive medium. As a result, the induced electric field distribution is primarily determined by tissue conductivity and coil geometry.

### Detailed Coil Alignment Strategy

This section provides a detailed description of the alignment strategies evaluated for TMS coil positioning. While the parcellation was not directly involved in the full body segmentation, it enables accurate positioning of the TMS coils to target the precentral motor cortex. The coil placement strategy used in this study differs from standard neuronavigated TMS practice, where coils are typically positioned tangentially to the scalp and oriented based on local cortical anatomy (e.g., targeting the hand knob). Instead, a simplified geometric alignment was adopted. This design was chosen to ensure a standardized and reproducible framework for comparing different coil geometries, rather than to replicate clinical coil orientation.

For circular coils, three alignment strategies were investigated: using the center of the coil (Figure S4a), as proposed by Deng et al. (2020), the mid-radius, which was equidistant from the inner and outer edges (Figure S4b), and the outer edge of the coil (Figure S4c). Each of these strategies was thoroughly tested.

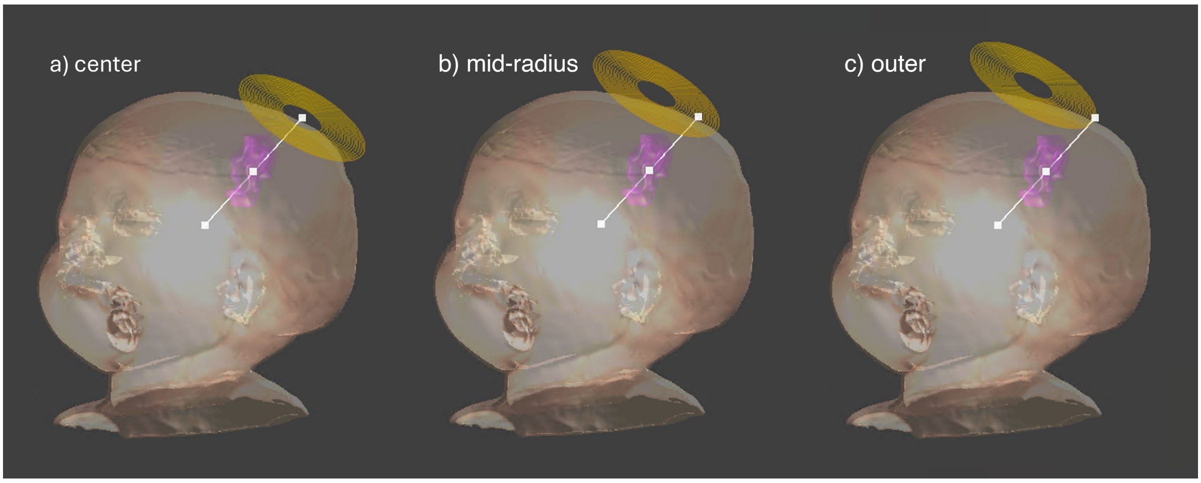


**Figure S4: Alignment Strategies for Circular Coil Positioning.** Screenshots from Sim4Life software illustrating the three alignment strategies employed for circular coils. The white line goes through the centroid of the skull, the centroid of the motor cortex parcellation and one of the three options: **(a)** alignment at center of the coil, **(b)** alignment at the mid-radius, which was equidistant from the inner and outer edges, and **(c)** alignment at the outer edge of the coil.

To determine the most suitable positioning among the three options, we compared the coordinates of the deepest point where the electric field magnitude reached $E_{\max}/2$ (Point D) of each coil. We used the coordinates of Coil 1 as a reference, as it was the smallest figure-of-eight coil (25mm) studied and thus expected to be the most precise. We calculated the distances between the deepest point of each coil and the point D reference.

This alignment strategy may influence the absolute electric field distributions, as more realistic tangential placements could modify field orientation and magnitude at the cortical surface. However, since all coils were evaluated under identical positioning constraints, the relative comparison between coil geometries is expected to remain robust.

### Spread-Based Analysis of TMS Coils

While the main manuscript focuses on electric field magnitude-based metrics, here we provide the detailed analysis of spatial spread and depth of stimulation, method inspired by Deng et al. (2020).

We quantified the spatial characteristics of stimulation by comparing $S_{1/2}$, the lateral spatial spread of regions exceeding $||\mathbf{E}_{\text{max}}||/2$, with $d_{1/2}$, the depth from the cortical surface to the deepest point at which the electric field remained above $||\mathbf{E}_{\text{max}}||/2$ (Figure S5). For each voxel where the field magnitude was at least $||\mathbf{E}_{\text{max}}||/2$, we calculated the minimum distance from that voxel to the cortical surface. The maximum of these minimum distances was denoted as $d_{1/2}$, representing the half-value depth. To quantify the spread of the electric field, we calculated the spread metric $S_{1/2}$ defined as:

$$S_{1/2} = \frac{V_{1/2}}{d_{1/2}} \tag{3}$$

The spread metric $S_{1/2}$ has units of $\text{kg/mm}^2$ and reflects the lateral spatial extent of stimulation relative to its depth. Lower values of $S_{1/2}$ indicate more focal electric fields.

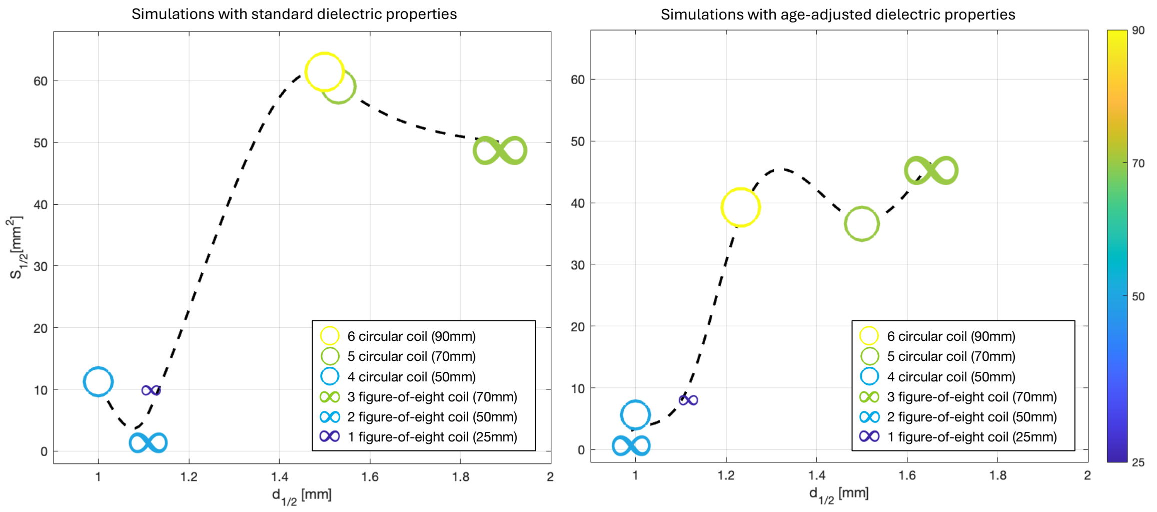


**Figure S5: Focality vs. Depth for TMS Coils.** Comparison of simulation results for different types of TMS coils. The two subplots display the relationship between the electric field focality, quantified by the half-value depth (d1/2) in mm (x-axis), and the half-value spread (S1/2) in $mm^2$ (y-axis). On the left, results are shown with standard dielectric properties; on the right, with age-adjusted dielectric properties. Both plots share the same axes and limits. The six types of coils compared are: (i) three circular coils of different sizes, represented by circles of varying sizes and colors corresponding to the coil size; (ii) three figure-of-eight coils, represented by infinity symbols of different sizes and colors according to the coil size. A smoothing spline is plotted to illustrate the general trend for the six coils in each subplot.

In evaluating the spread ($S_{1/2}$) on the cortical surface versus the depth ($d_{1/2}$) from the cortical surface to the point where the electric field exceeds $E_{max}/2$, larger coils demonstrate a trade-off between increased depth and spread. While larger coils offer deeper penetration, they also stimulate a larger volume, reducing focality. In contrast, smaller coils, such as the 25mm coil, produce a more localized and superficial electric field.

To determine the theoretically correct depth for a 10-month-old child, we considered that the cortical thickness of the precentral gyrus (motor cortex) in adults is approximately 3.34 mm (Biega et al., 2020). In comparison, the study by Tremblay et al. (2017) indicates that the precentral gyrus in a a 300-days-old child (10-month-old) has a thickness of approximately 2.60 mm. Layer IV in adults is located at a depth of about 1.30 to 1.60 mm (DeFelipe et al., 2002), suggesting that in a 10-month-old, this layer likely lies at a depth of approximately 1.01 to 1.25 mm. This depth range was defined as our target for effective stimulation. Given this information, it is noteworthy that the 50mm circular coil might not fully reach Layer IV. However, all other coils surpass the 1.01 mm depth, making them suitable for stimulating Layer IV.

Simulations performed with age-adjusted dielectric properties show similar trends. The distribution of coils in the plots and the trend curves consistently show similar patterns. Higher conductivity leads to an increase in $E_{max}$, which in turn raises $E_{max}/2$, ultimately reducing both the spread and depth, as shown in Figure S5.

#### Adult Electric Field Comparison

To go a bit deeper into our contextual discussion about the differences between adult and pediatric simulations, several methodological and biophysical factors should be considered.

Both studies operate under comparable simulation conditions, including similar excitation parameters (5 kA, 2.5 kHz) and the use of finite-element modeling within the Sim4Life environment. At this frequency range, electromagnetic interactions fall within the magnetoquasi-static regime, where induced electric fields are primarily governed by tissue conductivity.

One important difference lies in the dielectric properties used. Zhong et al. (2025) rely on the IT'IS database (v4.0), whereas our simulations use the updated IT'IS v4.1 dataset. The newer version incorporates revised low-frequency conductivity values based on more recent experimental measurements, generally leading to higher conductivity across multiple tissues (Hasgall et al., 2022). This increase can moderately enhance induced current flow and contribute to higher electric field magnitudes.

Differences in stimulation targets may also introduce variability. The adult study evaluates stimulation at the vertex and dorsolateral prefrontal cortex (DLPFC), whereas our simulations are aligned with the motor cortex. Although these regions differ functionally, they are located at comparable cortical depths, and therefore this factor is not expected to significantly affect global comparisons of Emax.

The dominant source of variation arises from anatomical and developmental differences. Pediatric subjects exhibit smaller head size, resulting in a reduced distance between the coil and cortical surface, which increases field strength due to reduced spatial decay. In addition, the skull in infants is thinner and presents higher effective conductivity, leading to reduced attenuation of the induced fields. These factors collectively enhance the transmission and concentration of electric fields in the brain.

However, this comparison remains indirect because the simulations were performed using different anatomical models and computational assumptions. Consequently, the observed differences in electric field magnitude should not be attributed solely to developmental anatomy but instead provide contextual comparison with previously published studies.

**Table S1: List of Segmented Head and Neck Tissues with Corresponding Volumes.** Tissues were grouped to extract data such as total cerebrospinal fluid (CSF) volume and brain parenchyma (functional part) volume, which were used to validate our model.

| **Tissue** | | | **Volume ($cm^3$)** |
|---|---|---|---|
| **Total CSF Brain** | | | **93.5654** |
| CSF_Brain | 68.1175 | Fourth_Ventricle | 2.5702 |
| Lateral_Ventricle_L | 11.4675 | Third_Ventricle | 0.9772 |
| Lateral_Ventricle_R | 10.4331 | | |
| **Brain_Parenchyma** | | | **859.8109** |
| Amygdala_L | 0.8536 | Hippocampus_R | 2.7909 |
| Amygdala_R | 0.9285 | Hypothalamus_L | 0.3422 |
| Brain_Gray_Matter_L | 252.9300 | Hypothalamus_R | 0.3491 |
| Brain_Gray_Matter_R | 263.9070 | Mammillary_Bodies | 0.0865 |
| Brain_White_Matter_L | 104.2740 | Medulla_Oblongata | 2.1037 |
| Brain_White_Matter_R | 108.4650 | Midbrain | 2.1983 |
| Caudate_L | 3.3035 | Pituitary | 0.1813 |
| Caudate_R | 3.1618 | Pons | 5.2099 |
| Cerebellum_Gray_Matter_L | 25.4160 | Putamen_L | 4.5616 |
| Cerebellum_Gray_Matter_R | 24.0938 | Putamen_R | 4.6533 |
| Cerebellum_White_Matter_L | 9.3961 | Septum_Pellucidum | 0.2684 |
| Cerebellum_White_Matter_R | 9.6973 | Thalamus_L | 7.1336 |
| Choroid_Plexus | 0.9859 | Thalamus_R | 6.7068 |
| Colliculi | 0.3025 | Ventral_Diencephalon_L | 1.4953 |
| Globus_Pallidus_L | 1.9807 | Ventral_Diencephalon_R | 1.6510 |
| Globus_Pallidus_R | 1.8571 | Vermis_Gray_Matter | 3.7249 |
| Hippocampus_L | 2.7294 | Vermis_White_Matter | 2.0763 |
| **Eyes/Optic** | | | **32.6097** |
| Extraocular_Muscles | 3.4293 | Eye_Sclera_L | 0.7394 |
| Eye_Sclera_R | 0.7101 | Eye_Choroid_L | 0.6181 |
| Eye_Choroid_R | 0.6126 | Eye_Vitreus_L | 2.0122 |
| Eye_Vitreus_R | 2.0136 | Eye_Ciliary_Muscle_L | 0.0363 |
| Eye_Ciliary_Muscle_R | 0.0307 | Intraorbital_Fat | 19.7351 |
| Eye_Cornea_L | 0.1317 | Eye_Cornea_R | 0.1120 |
| Eye_Lens_L | 0.2429 | Eye_Lens_R | 0.2034 |
| Eye_Retina_L | 0.5124 | Eye_Retina_R | 0.5155 |
| Optic_Chiasm | 0.2850 | Optic_Nerve_L | 0.1734 |
| Optic_Nerve_R | 0.2103 | Optic_Tract_L | 0.1348 |
| Optic_Tract_R | 0.1524 | | |
| **Head & Neck** | | | **468.6377** |
| Air_Head_Neck | 20.5436 | Salivary_Glands | 15.7722 |
| Esophagus | 1.7458 | Skull_BM | 27.6315 |
| Lower_Mandible_BM | 4.5070 | Skull_CB | 206.952 |
| Lower_Mandible_CB | 13.1153 | Teeth | 3.73275 |
| Meniges_Brain | 149.2250 | Tongue | 14.0108 |
| Mucosa | 4.6369 | Trachea | 1.9077 |
| Nasal_Cartillage | 2.7399 | Turbinates | 2.1172 |
| **Full Body** | | | **6510.3969** |
| Arteries_Head_Neck | 7.9631 | Skin | 642.7280 |
| Connective_Tissue_Body | 821.5430 | Subcutaneous_Fat_Body | 2721.5700 |
| Connective_Tissue_Head_Neck | 5.7441 | Subcutaneous_Fat_Head_Neck | 284.2760 |
| Muscle_Body | 1791.2400 | Vessels_Body | 51.6970 |
| Muscle_Head_Neck | 170.6510 | Vein_Head_Neck | 12.9847 |
| **Visceral Tissues** | | | **985.9498** |
| Adrenal_L | 0.1640 | Ovaries | 0.5130 |
| Adrenal_R | 0.0764 | Pancreas | 7.8496 |
| Fallopian_Tubes | 0.1030 | Small_Bowel_Air | 59.4959 |
| Gall_Bladder | 0.6835 | Small_Bowel_Air_Wall | 98.6431 |

*Continued on next page*

| | | | |
|---|---|---|---|
| Gallbladder_Wall | 1.8416 | Small_Bowel_Contents | 18.6867 |
| Heart | 44.7026 | Spleen | 27.8337 |
| Inta_Abdominal_Fat | 42.3634 | Stomach_Contents_Air | 1.6871 |
| Kidney_L | 20.3153 | Stomach_Contents_Solid | 5.3147 |
| Kidney_R | 25.1170 | Stomach_Wall | 5.9857 |
| Large_Bowel_Contents_Air | 40.9016 | Thymus | 21.4687 |
| Large_Bowel_Contents_Solid | 44.0668 | Urinary_Bladder | 18.0844 |
| Large_Bowel_Wall | 43.2489 | Urinary_Bladder_Wall | 10.5614 |
| Liver_Minus_Vessels | 238.7400 | Uterus | 0.7035 |
| Lung_L | 97.3817 | Vagina | 2.2735 |
| Lung_R | 107.1430 | | |
| **Spine** | | | **153.6294** |
| Coccyx_CB | 0.0506 | Coccyx_BM | 0.0023 |
| Vertebra_C1_CB | 1.2738 | Vertebra_C1_BM | 0.0763 |
| Vertebra_C2_CB | 1.7708 | Vertebra_C2_BM | 0.7422 |
| Vertebra_C3_CB | 1.1790 | Vertebra_C3_BM | 0.1897 |
| Vertebra_C4_CB | 1.1742 | Vertebra_C4_BM | 0.1989 |
| Vertebra_C5_CB | 0.8257 | Vertebra_C5_BM | 0.1635 |
| Vertebra_C6_CB | 0.9125 | Vertebra_C6_BM | 0.2010 |
| Vertebra_C7_CB | 0.8918 | Vertebra_C7_BM | 0.2002 |
| Vertebra_T1_CB | 0.8676 | Vertebra_T1_BM | 0.1973 |
| Vertebra_T2_CB | 0.9182 | Vertebra_T2_BM | 0.3397 |
| Vertebra_T3_CB | 1.0499 | Vertebra_T3_BM | 0.4351 |
| Vertebra_T4_CB | 1.4094 | Vertebra_T4_BM | 0.6077 |
| Vertebra_T5_CB | 1.2331 | Vertebra_T5_BM | 0.5200 |
| Vertebra_T6_CB | 1.1622 | Vertebra_T6_BM | 0.4605 |
| Vertebra_T7_CB | 1.3603 | Vertebra_T7_BM | 0.5696 |
| Vertebra_T8_CB | 1.3388 | Vertebra_T8_BM | 0.6098 |
| Vertebra_T9_CB | 1.5608 | Vertebra_T9_BM | 0.6535 |
| Vertebra_T10_CB | 1.5626 | Vertebra_T10_BM | 0.7987 |
| Vertebra_T11_CB | 1.5475 | Vertebra_T11_BM | 0.7884 |
| Vertebra_T12_CB | 1.5720 | Vertebra_T12_BM | 0.7384 |
| Vertebra_L1_CB | 1.7056 | Vertebra_L1_BM | 0.9043 |
| Vertebra_L2_CB | 1.6029 | Vertebra_L2_BM | 1.0155 |
| Vertebra_L3_CB | 1.7483 | Vertebra_L3_BM | 1.2020 |
| Vertebra_L4_CB | 1.6958 | Vertebra_L4_BM | 1.0069 |
| Vertebra_L5_CB | 1.5782 | Vertebra_L5_BM | 0.9497 |
| Vertebra_S1_CB | 3.8371 | Vertebra_S1_BM | 0.9217 |
| Vertebra_S2_CB | 2.5166 | Vertebra_S2_BM | 0.4058 |
| Vertebra_S3_CB | 1.7916 | Vertebra_S3_BM | 0.4651 |
| Vertebra_S4_CB | 0.6067 | Vertebra_S4_BM | 0.0404 |
| Vertebra_S5_CB | 0.1532 | Vertebra_S5_BM | 0.0418 |
| Cauda_Equina | 0.5408 | Connective_Tissues_Spine | 0.6303 |
| CSF_Spine | 19.8892 | Intervertebral_Discs | 17.3490 |
| Ligaments_Spine | 51.7717 | Meniges_Spine | 6.3520 |
| Spinal_Cord | 5.4391 | | |
| **Thoracic cage** | | | **46.9636** |
| Rib_T1_L_CB | 0.7271 | Rib_T1_L_BM | 0.0772 |
| Rib_T1_R_CB | 0.6314 | Rib_T1_R_BM | 0.0794 |
| Rib_T2_L_CB | 0.9047 | Rib_T2_L_BM | 0.0767 |
| Rib_T2_R_CB | 0.6812 | Rib_T2_R_BM | 0.0350 |
| Rib_T3_L_CB | 1.0647 | Rib_T3_L_BM | 0.0776 |
| Rib_T3_R_CB | 0.9892 | Rib_T3_R_BM | 0.0801 |
| Rib_T4_L_CB | 1.4137 | Rib_T4_L_BM | 0.2780 |
| Rib_T4_R_CB | 1.2996 | Rib_T4_R_BM | 0.2215 |
| Rib_T5_L_CB | 1.5759 | Rib_T5_L_BM | 0.3711 |

*Continued on next page*

| | | | |
|---|---|---|---|
| Rib_T5_R_CB | 1.4778 | Rib_T5_R_BM | 0.3635 |
| Rib_T6_L_CB | 1.6274 | Rib_T6_L_BM | 0.3901 |
| Rib_T6_R_CB | 1.6114 | Rib_T6_R_BM | 0.3929 |
| Rib_T7_L_CB | 1.6705 | Rib_T7_L_BM | 0.2818 |
| Rib_T7_R_CB | 1.6484 | Rib_T7_R_BM | 0.3642 |
| Rib_T8_L_CB | 2.0427 | Rib_T8_L_BM | 0.5818 |
| Rib_T8_R_CB | 1.9061 | Rib_T8_R_BM | 0.5068 |
| Rib_T9_L_CB | 2.1714 | Rib_T9_L_BM | 0.5976 |
| Rib_T9_R_CB | 2.0973 | Rib_T9_R_BM | 0.6021 |
| Rib_T10_L_CB | 2.0790 | Rib_T10_L_BM | 0.5591 |
| Rib_T10_R_CB | 2.0083 | Rib_T10_R_BM | 0.5935 |
| Rib_T11_L_CB | 1.0511 | Rib_T11_L_BM | 0.1758 |
| Rib_T11_R_CB | 1.1841 | Rib_T11_R_BM | 0.3202 |
| Rib_T12_L_CB | 0.5423 | Rib_T12_L_BM | 0.0892 |
| Rib_T12_R_CB | 0.6125 | Rib_T12_R_BM | 0.1767 |
| Sternum_CB | 3.1561 | Sternum_BM | 3.4978 |
| **Bones Upper Limb** | | | **62.6573** |
| Clavicle_L_CB | 1.4850 | Clavicle_L_BM | 1.1555 |
| Clavicle_R_CB | 1.4793 | Clavicle_R_BM | 1.0602 |
| Humerus_L_CB | 4.3540 | Humerus_L_BM | 8.6561 |
| Humerus_R_CB | 4.2409 | Humerus_R_BM | 8.6391 |
| Radius_L_CB | 2.2447 | Radius_L_BM | 2.3260 |
| Radius_R_CB | 2.1603 | Radius_R_BM | 2.1344 |
| Scapula_L_CB | 3.4237 | Scapula_L_BM | 3.9940 |
| Scapula_R_CB | 3.6856 | Scapula_R_BM | 4.3105 |
| Ulna_L_CB | 1.8928 | Ulna_L_BM | 1.6602 |
| Ulna_R_CB | 2.0454 | Ulna_R_BM | 1.7097 |
| **Bones Hand** | | | **9.1917** |
| 1st_Distal_Phalanx_L_CB | 0.0532 | 1st_Distal_Phalanx_L_BM | 0.0108 |
| 1st_Distal_Phalanx_R_CB | 0.0634 | 1st_Distal_Phalanx_R_BM | 0.0133 |
| 1st_Metacarpal_L_CB | 0.1966 | 1st_Metacarpal_L_BM | 0.0863 |
| 1st_Metacarpal_R_CB | 0.1644 | 1st_Metacarpal_R_BM | 0.0627 |
| 1st_Proximal_Phalanx_L_CB | 0.2308 | 1st_Proximal_Phalanx_L_BM | 0.0881 |
| 1st_Proximal_Phalanx_R_CB | 0.1525 | 1st_Proximal_Phalanx_R_BM | 0.0425 |
| 2nd_Distal_Phalanx_L_CB | 0.0400 | 2nd_Distal_Phalanx_L_BM | 0.0061 |
| 2nd_Distal_Phalanx_R_CB | 0.0526 | 2nd_Distal_Phalanx_R_BM | 0.0089 |
| 2nd_Metacarpal_L_CB | 0.4236 | 2nd_Metacarpal_L_BM | 0.2206 |
| 2nd_Metacarpal_R_CB | 0.3265 | 2nd_Metacarpal_R_BM | 0.1288 |
| 2nd_Mid_Phalanx_L_CB | 0.1154 | 2nd_Mid_Phalanx_L_BM | 0.0375 |
| 2nd_Mid_Phalanx_R_CB | 0.0702 | 2nd_Mid_Phalanx_R_BM | 0.0115 |
| 2nd_Proximal_Phalanx_L_CB | 0.1822 | 2nd_Proximal_Phalanx_L_BM | 0.0809 |
| 2nd_Proximal_Phalanx_R_CB | 0.1523 | 2nd_Proximal_Phalanx_R_BM | 0.0636 |
| 3rd_Distal_Phalanx_L_CB | 0.0933 | 3rd_Distal_Phalanx_L_BM | 0.0246 |
| 3rd_Distal_Phalanx_R_CB | 0.0699 | 3rd_Distal_Phalanx_R_BM | 0.0150 |
| 3rd_Metacarpal_L_CB | 0.2850 | 3rd_Metacarpal_L_BM | 0.1076 |
| 3rd_Metacarpal_R_CB | 0.4051 | 3rd_Metacarpal_R_BM | 0.1892 |
| 3rd_Mid_Phalanx_L_CB | 0.2077 | 3rd_Mid_Phalanx_L_BM | 0.0952 |
| 3rd_Mid_Phalanx_R_CB | 0.0634 | 3rd_Mid_Phalanx_R_BM | 0.0143 |
| 3rd_Proximal_Phalanx_L_CB | 0.1836 | 3rd_Proximal_Phalanx_L_BM | 0.0729 |
| 3rd_Proximal_Phalanx_R_CB | 0.1457 | 3rd_Proximal_Phalanx_R_BM | 0.0586 |
| 4th_Distal_Phalanx_L_CB | 0.0546 | 4th_Distal_Phalanx_L_BM | 0.0108 |
| 4th_Distal_Phalanx_R_CB | 0.0503 | 4th_Distal_Phalanx_R_BM | 0.0083 |
| 4th_Metacarpal_L_CB | 0.2433 | 4th_Metacarpal_L_BM | 0.1008 |
| 4th_Metacarpal_R_CB | 0.3184 | 4th_Metacarpal_R_BM | 0.1446 |
| 4th_Mid_Phalanx_L_CB | 0.1312 | 4th_Mid_Phalanx_L_BM | 0.0343 |
| 4th_Mid_Phalanx_R_CB | 0.0739 | 4th_Mid_Phalanx_R_BM | 0.0171 |

*Continued on next page*

| | | | |
|---|---|---|---|
| 4th_Proximal_Phalanx_L_CB | 0.4709 | 4th_Proximal_Phalanx_L_BM | 0.0538 |
| 4th_Proximal_Phalanx_R_CB | 0.1581 | 4th_Proximal_Phalanx_R_BM | 0.0452 |
| 5th_Distal_Phalanx_L_CB | 0.0472 | 5th_Distal_Phalanx_L_BM | 0.0071 |
| 5th_Distal_Phalanx_R_CB | 0.0429 | 5th_Distal_Phalanx_R_BM | 0.0057 |
| 5th_Metacarpal_L_CB | 0.2899 | 5th_Metacarpal_L_BM | 0.1283 |
| 5th_Metacarpal_R_CB | 0.2561 | 5th_Metacarpal_R_BM | 0.1033 |
| 5th_Mid_Phalanx_L_CB | 0.0756 | 5th_Mid_Phalanx_L_BM | 0.0153 |
| 5th_Mid_Phalanx_R_CB | 0.0627 | 5th_Mid_Phalanx_R_BM | 0.0143 |
| 5th_Proximal_Phalanx_L_CB | 0.0633 | 5th_Proximal_Phalanx_L_BM | 0.0137 |
| 5th_Proximal_Phalanx_R_CB | 0.1939 | 5th_Proximal_Phalanx_R_BM | 0.0887 |
| Carpal_Bones_L_CB | 0.1816 | Carpal_Bones_L_BM | 0.0479 |
| Carpal_Bones_R_CB | 0.3355 | Carpal_Bones_R_BM | 0.1863 |
| **Bones Lower Limb** | | | **157.2671** |
| Calcaneus_L_CB | 1.4246 | Calcaneus_L_BM | 2.4549 |
| Calcaneus_R_CB | 1.3128 | Calcaneus_R_BM | 2.1986 |
| Femur_L_CB | 7.6599 | Femur_L_BM | 21.7107 |
| Femur_R_CB | 8.0487 | Femur_R_BM | 22.8400 |
| Fibula_L_CB | 1.9063 | Fibula_L_BM | 1.3671 |
| Fibula_R_CB | 2.0764 | Fibula_R_BM | 1.5169 |
| Iliac_Bone_L_CB | 6.7293 | Iliac_Bone_L_BM | 12.2222 |
| Iliac_Bone_R_CB | 6.8290 | Iliac_Bone_R_BM | 13.1659 |
| Kneecap_L_CB | 0.8553 | Kneecap_L_BM | 1.4413 |
| Kneecap_R_CB | 0.7089 | Kneecap_R_BM | 0.9422 |
| Talus_L_CB | 0.9878 | Talus_L_BM | 1.4919 |
| Talus_R_CB | 0.8301 | Talus_R_BM | 1.1667 |
| Tibia_L_CB | 5.6516 | Tibia_L_BM | 12.5131 |
| Tibia_R_CB | 5.3333 | Tibia_R_BM | 11.8814 |
| **Bones Feet** | | | **11.5330** |
| 1st_Distal_Phalanx_L_CB | 0.0730 | 1st_Distal_Phalanx_L_BM | 0.0231 |
| 1st_Distal_Phalanx_R_CB | 0.0951 | 1st_Distal_Phalanx_R_BM | 0.0288 |
| 1st_Metatarsal_L_CB | 0.5816 | 1st_Metatarsal_L_BM | 0.4925 |
| 1st_Metatarsal_R_CB | 0.7098 | 1st_Metatarsal_R_BM | 0.6130 |
| 1st_Prox_Phalanx_L_CB | 0.2618 | 1st_Prox_Phalanx_L_BM | 0.0858 |
| 1st_Prox_Phalanx_R_CB | 0.2796 | 1st_Prox_Phalanx_R_BM | 0.1806 |
| 2nd_Distal_Phalanx_L_CB | 0.0316 | 2nd_Distal_Phalanx_L_BM | 0.0060 |
| 2nd_Distal_Phalanx_R_CB | 0.0328 | 2nd_Distal_Phalanx_R_BM | 0.0042 |
| 2nd_Metatarsal_L_CB | 0.4335 | 2nd_Metatarsal_L_BM | 0.2140 |
| 2nd_Metatarsal_R_CB | 0.4658 | 2nd_Metatarsal_R_BM | 0.2566 |
| 2nd_Mid_Phalanx_L_CB | 0.0484 | 2nd_Mid_Phalanx_L_BM | 0.0117 |
| 2nd_Mid_Phalanx_R_CB | 0.0410 | 2nd_Mid_Phalanx_R_BM | 0.0072 |
| 2nd_Prox_Phalanx_L_CB | 0.0814 | 2nd_Prox_Phalanx_L_BM | 0.0228 |
| 2nd_Prox_Phalanx_R_CB | 0.1008 | 2nd_Prox_Phalanx_R_BM | 0.0330 |
| 3rd_Distal_Phalanx_L_CB | 0.0253 | 3rd_Distal_Phalanx_L_BM | 0.0025 |
| 3rd_Distal_Phalanx_R_CB | 0.0221 | 3rd_Distal_Phalanx_R_BM | 0.0017 |
| 3rd_Metatarsal_L_CB | 0.2805 | 3rd_Metatarsal_L_BM | 0.1170 |
| 3rd_Metatarsal_R_CB | 0.4475 | 3rd_Metatarsal_R_BM | 0.2413 |
| 3rd_Mid_Phalanx_L_CB | 0.0289 | 3rd_Mid_Phalanx_L_BM | 0.0042 |
| 3rd_Mid_Phalanx_R_CB | 0.0384 | 3rd_Mid_Phalanx_R_BM | 0.0057 |
| 3rd_Prox_Phalanx_L_CB | 0.0529 | 3rd_Prox_Phalanx_L_BM | 0.0089 |
| 3rd_Prox_Phalanx_R_CB | 0.0696 | 3rd_Prox_Phalanx_R_BM | 0.0149 |
| 4th_Distal_Phalanx_L_CB | 0.0229 | 4th_Distal_Phalanx_L_BM | 0.0021 |
| 4th_Distal_Phalanx_R_CB | 0.0215 | 4th_Distal_Phalanx_R_BM | 0.0014 |
| 4th_Metatarsal_L_CB | 0.2321 | 4th_Metatarsal_L_BM | 0.0678 |
| 4th_Metatarsal_R_CB | 0.4084 | 4th_Metatarsal_R_BM | 0.2146 |
| 4th_Mid_Phalanx_L_CB | 0.0308 | 4th_Mid_Phalanx_L_BM | 0.0037 |
| 4th_Mid_Phalanx_R_CB | 0.0328 | 4th_Mid_Phalanx_R_BM | 0.0043 |

*Continued on next page*

| | | | |
|---|---:|---|---:|
| 4th_Prox_Phalanx_L_CB | 0.0482 | 4th_Prox_Phalanx_L_BM | 0.0073 |
| 4th_Prox_Phalanx_R_CB | 0.0730 | 4th_Prox_Phalanx_R_BM | 0.01595 |
| 5th_Distal_Phalanx_L_CB | 0.0178 | 5th_Distal_Phalanx_L_BM | 0.0011 |
| 5th_Distal_Phalanx_R_CB | 0.0180 | 5th_Distal_Phalanx_R_BM | 0.0007 |
| 5th_Metatarsal_L_CB | 0.3493 | 5th_Metatarsal_L_BM | 0.1499 |
| 5th_Metatarsal_R_CB | 0.4158 | 5th_Metatarsal_R_BM | 0.2094 |
| 5th_Mid_Phalanx_L_CB | 0.0280 | 5th_Mid_Phalanx_L_BM | 0.0025 |
| 5th_Mid_Phalanx_R_CB | 0.0264 | 5th_Mid_Phalanx_R_BM | 0.0018 |
| 5th_Prox_Phalanx_L_CB | 0.0735 | 5th_Prox_Phalanx_L_BM | 0.0165 |
| 5th_Prox_Phalanx_R_CB | 0.0707 | 5th_Prox_Phalanx_R_BM | 0.0159 |
| Cuboid_L_CB | 0.4489 | Cuboid_L_BM | 0.4788 |
| Cuboid_R_CB | 0.5258 | Cuboid_R_BM | 0.6032 |
| Cuneiform_L_CB | 0.0821 | Cuneiform_L_BM | 0.0411 |
| Cuneiform_R_CB | 0.1224 | Cuneiform_R_BM | 0.0701 |

**Table S2: Dielectric Properties and Age-Dependant Adjustments for Various Tissues.**

| Tissue | Assigned Material | Perm. at 2.5kHz | Perm. Ratio | Perm. Adjusted | Cond. at 2.5kHz | Cond. Ratio | Cond. Adjusted |
|---|---|---|---|---|---|---|---|
| Air_Head_Neck | Air | 1.00 | 1.00 | 1.00 | 0.0000 | 1.00 | 0.0000 |
| Amygdala_L | Brain (Gray Matter) | 78103.96 | 1.43 | 111800.48 | 0.1043 | 1.65 | 0.1721 |
| Amygdala_R | Brain (Gray Matter) | 78103.96 | 1.43 | 111800.48 | 0.1043 | 1.65 | 0.1721 |
| Arteries_Head_Neck | Blood | 5256.78 | 1.00 | 5256.78 | 0.7000 | 1.00 | 0.7000 |
| Brain_Gray_Matter_L | Brain (Gray Matter) | 78103.96 | 1.43 | 111800.48 | 0.1043 | 1.65 | 0.1721 |
| Brain_Gray_Matter_R | Brain (Gray Matter) | 78103.96 | 1.43 | 111800.48 | 0.1043 | 1.65 | 0.1721 |
| Brain_White_Matter_L | Brain (White Matter) | 34282.00 | 1.43 | 49072.34 | 0.0645 | 1.65 | 0.1065 |
| Brain_White_Matter_R | Brain (White Matter) | 34282.00 | 1.43 | 49072.34 | 0.0645 | 1.65 | 0.1065 |
| Caudate_L | Brain (Gray Matter) | 78103.96 | 1.43 | 111800.48 | 0.1043 | 1.65 | 0.1721 |
| Caudate_R | Brain (Gray Matter) | 78103.96 | 1.43 | 111800.48 | 0.1043 | 1.65 | 0.1721 |
| Cerebellum_Gray_Matter_L | Cerebellum | 78398.90 | 1.43 | 112222.67 | 0.1243 | 1.65 | 0.2051 |
| Cerebellum_Gray_Matter_R | Cerebellum | 78398.90 | 1.43 | 112222.67 | 0.1243 | 1.65 | 0.2051 |
| Cerebellum_White_Matter_L | Brain (White Matter) | 34282.00 | 1.43 | 49072.34 | 0.0645 | 1.65 | 0.1065 |
| Cerebellum_White_Matter_R | Brain (White Matter) | 34282.00 | 1.43 | 49072.34 | 0.0645 | 1.65 | 0.1065 |
| Choroid_Plexus | Blood | 5256.78 | 1.00 | 5256.78 | 0.7000 | 1.00 | 0.7000 |
| Colliculi | Brain (Gray Matter) | 78103.96 | 1.43 | 111800.48 | 0.1043 | 1.65 | 0.1721 |
| CSF_Brain | Cerebrospinal Fluid | 109.00 | 1.00 | 109.00 | 2.0000 | 1.00 | 2.0000 |
| CSF_Spine | Cerebrospinal Fluid | 109.00 | 1.00 | 109.00 | 2.0000 | 1.00 | 2.0000 |
| Extraocular_Muscles | Muscle | 124369.75 | 1.36 | 169451.67 | 0.3316 | 1.62 | 0.5360 |
| Eye_Choroid_L | Blood | 5256.78 | 1.00 | 5256.78 | 0.7000 | 1.00 | 0.7000 |
| Eye_Choroid_R | Blood | 5256.78 | 1.00 | 5256.78 | 0.7000 | 1.00 | 0.7000 |
| Eye_Ciliary_Muscle_L | Muscles | 124369.75 | 1.36 | 169451.67 | 0.3316 | 1.62 | 0.5360 |
| Eye_Ciliary_Muscle_R | Muscles | 124369.75 | 1.36 | 169451.67 | 0.3316 | 1.62 | 0.5360 |
| Eye_Cornea_L | Eye (Cornea) | 88636.15 | 1.32 | 117074.47 | 0.4256 | 1.55 | 0.6594 |
| Eye_Cornea_R | Eye (Cornea) | 88636.15 | 1.32 | 117074.47 | 0.4256 | 1.55 | 0.6594 |
| Eye_Lens_L | Eye (Lens) | 974.50 | 1.32 | 1287.17 | 0.2000 | 1.55 | 0.3099 |
| Eye_Lens_R | Eye (Lens) | 974.50 | 1.32 | 1287.17 | 0.2000 | 1.55 | 0.3099 |
| Eye_Retina_L | Brain (Gray Matter) | 78103.96 | 1.43 | 111800.48 | 0.1043 | 1.65 | 0.1721 |
| Eye_Retina_R | Brain (Gray Matter) | 78103.96 | 1.43 | 111800.48 | 0.1043 | 1.65 | 0.1721 |
| Eye_Sclera_L | Eye (Sclera) | 28958.11 | 1.32 | 38249.13 | 0.5069 | 1.55 | 0.7854 |
| Eye_Sclera_R | Eye (Sclera) | 28958.11 | 1.32 | 38249.13 | 0.5069 | 1.55 | 0.7854 |
| Eye_Vitreus_L | Eye (Vitrous Humor) | 98.98 | 1.00 | 98.98 | 1.5000 | 1.00 | 1.5000 |
| Eye_Vitreus_R | Eye (Vitrous Humor) | 98.98 | 1.00 | 98.98 | 1.5000 | 1.00 | 1.5000 |

*Continued on next page*

| **Tissue** | **Assigned Material** | **Perm. at 2.5kHz** | **Perm. Ratio** | **Perm. Adjusted** | **Cond. at 2.5kHz** | **Cond. Ratio** | **Cond. Adjusted** |
|---|---|---|---|---|---|---|---|
| Fourth_Ventricle | Cerebrospinal Fluid | 109.00 | 1.00 | 109.00 | 2.0000 | 1.00 | 2.0000 |
| Globus_Pallidus_L | Brain (Gray Matter) | 78103.96 | 1.43 | 111800.48 | 0.1043 | 1.65 | 0.1721 |
| Globus_Pallidus_R | Brain (Gray Matter) | 78103.96 | 1.43 | 111800.48 | 0.1043 | 1.65 | 0.1721 |
| Hippocampus_L | Brain (Gray Matter) | 78103.96 | 1.43 | 111800.48 | 0.1043 | 1.65 | 0.1721 |
| Hippocampus_R | Brain (Gray Matter) | 78103.96 | 1.43 | 111800.48 | 0.1043 | 1.65 | 0.1721 |
| Hypophysis | Thyroid gland | 28024.24 | 1.34 | 37586.02 | 0.3850 | 1.60 | 0.6172 |
| Hypothalamus_L | Brain (Gray Matter) | 78103.96 | 1.43 | 111800.48 | 0.1043 | 1.65 | 0.1721 |
| Hypothalamus_R | Brain (Gray Matter) | 78103.96 | 1.43 | 111800.48 | 0.1043 | 1.65 | 0.1721 |
| Intervertebral_Discs | Intervertebral Disc | 60.69 | 1.32 | 80.17 | 0.8300 | 1.55 | 1.2860 |
| Intraorbital_Fat | Fat (Average Infiltrated) | 6425.32 | 1.32 | 8486.84 | 0.0424 | 1.55 | 0.0657 |
| Lateral_Ventricle_L | Cerebrospinal Fluid | 109.00 | 1.00 | 109.00 | 2.0000 | 1.00 | 2.0000 |
| Lateral_Ventricle_R | Cerebrospinal Fluid | 109.00 | 1.00 | 109.00 | 2.0000 | 1.00 | 2.0000 |
| Ligaments_Spine | Tendon/Ligament | 54469.28 | 1.32 | 71945.39 | 0.3850 | 1.55 | 0.5965 |
| Lower_Mandible_BM | Bone Marrow (Red) | 2471.32 | 1.32 | 3264.23 | 0.1020 | 1.55 | 0.1580 |
| Lower_Mandible_CB | Bone (Cortical) | 1435.18 | 2.04 | 2923.94 | 0.0203 | 3.50 | 0.0709 |
| Mammilary_Bodies | Brain (Gray Matter) | 78103.96 | 1.43 | 111800.48 | 0.1043 | 1.65 | 0.1721 |
| Medulla_Oblongata | Cerebellum | 78398.90 | 1.43 | 112222.67 | 0.1243 | 1.65 | 0.2051 |
| Meniges_Brain | Dura | 2742.69 | 1.43 | 3925.98 | 0.5010 | 1.65 | 0.8264 |
| Meniges_Spine | Dura | 2742.69 | 1.43 | 3925.98 | 0.5010 | 1.65 | 0.8264 |
| Midbrain | Cerebellum | 78398.90 | 1.43 | 112222.67 | 0.1243 | 1.65 | 0.2051 |
| Mucosa | Muscles | 124369.75 | 1.36 | 169451.67 | 0.3316 | 1.62 | 0.5360 |
| Muscle_Head_Neck | Muscles | 124369.75 | 1.36 | 169451.67 | 0.3316 | 1.62 | 0.5360 |
| Nasal_Cartillage | Cartilage | 11324.33 | 1.32 | 14957.66 | 0.1751 | 1.55 | 0.2713 |
| Optic_Chiasm | Nerve | 59931.17 | 1.32 | 79159.69 | 0.0306 | 1.55 | 0.0474 |
| Optic_Nerve_L | Nerve | 59931.17 | 1.32 | 79159.69 | 0.0306 | 1.55 | 0.0474 |
| Optic_Nerve_R | Nerve | 59931.17 | 1.32 | 79159.69 | 0.0306 | 1.55 | 0.0474 |
| Optic_Tract_L | Nerve | 59931.17 | 1.32 | 79159.69 | 0.0306 | 1.55 | 0.0474 |
| Optic_Tract_R | Nerve | 59931.17 | 1.32 | 79159.69 | 0.0306 | 1.55 | 0.0474 |
| Pons | Cerebellum | 78398.90 | 1.43 | 112222.67 | 0.1243 | 1.65 | 0.2051 |
| Putamen_L | Brain (Gray Matter) | 78103.96 | 1.43 | 111800.48 | 0.1043 | 1.65 | 0.1721 |
| Putamen_R | Brain (Gray Matter) | 78103.96 | 1.43 | 111800.48 | 0.1043 | 1.65 | 0.1721 |
| Salivary_Glands | Salivary Gland | 136.80 | 1.34 | 183.48 | 0.6700 | 1.60 | 1.0741 |
| Septum_Pellucidum | Brain (White Matter) | 34282.00 | 1.43 | 49072.34 | 0.0645 | 1.65 | 0.1065 |
| Skin_Head_Neck | Skin (Dry) | 1135.21 | 1.67 | 1900.38 | 0.0002 | 1.94 | 0.0004 |
| Skull_BM | Bone Marrow (Red) | 2471.32 | 1.32 | 3264.23 | 0.1020 | 1.55 | 0.1580 |

*Continued on next page*

| Tissue | Assigned Material | Perm. at 2.5kHz | Perm. Ratio | Perm. Adjusted | Cond. at 2.5kHz | Cond. Ratio | Cond. Adjusted |
|---|---|---|---|---|---|---|---|
| Skull_CB | Bone (Cortical) | 1435.18 | 2.04 | 2923.94 | 0.0203 | 3.50 | 0.0709 |
| Spinal_Cord | Nerve | 59931.17 | 1.32 | 79159.69 | 0.0306 | 1.55 | 0.0474 |
| Fat_Head_Neck | Fat (Average Infiltrated) | 6425.32 | 1.32 | 8486.84 | 0.0424 | 1.55 | 0.0657 |
| Teeth | Bone (Cortical) | 1435.18 | 2.04 | 2923.94 | 0.0203 | 3.50 | 0.0709 |
| Thalamus_L | Brain (Gray Matter) | 78103.96 | 1.43 | 111800.48 | 0.1043 | 1.65 | 0.1721 |
| Thalamus_R | Brain (Gray Matter) | 78103.96 | 1.43 | 111800.48 | 0.1043 | 1.65 | 0.1721 |
| Third_Ventricle | Cerebrospinal Fluid | 109.00 | 1.00 | 109.00 | 2.0000 | 1.00 | 2.0000 |
| Tongue | Tongue | 29518.14 | 1.36 | 40217.96 | 0.2764 | 1.62 | 0.4468 |
| Turbinates | Muscles | 124369.75 | 1.36 | 169451.67 | 0.3316 | 1.62 | 0.5360 |
| Vein_Head_Neck | Blood | 5256.78 | 1.00 | 5256.78 | 0.7000 | 1.00 | 0.7000 |
| Ventral_Diencephalon_L | Brain (Gray Matter) | 78103.96 | 1.43 | 111800.48 | 0.1043 | 1.65 | 0.1721 |
| Ventral_Diencephalon_R | Brain (Gray Matter) | 78103.96 | 1.43 | 111800.48 | 0.1043 | 1.65 | 0.1721 |
| Vermis_Gray_Matter | Cerebellum | 78398.90 | 1.43 | 112222.67 | 0.1243 | 1.65 | 0.2051 |
| Vermis_White_Matter | Brain (White Matter) | 34282.00 | 1.43 | 49072.34 | 0.0645 | 1.65 | 0.1065 |
| Vertebra_C1_BM | Bone Marrow (Red) | 2471.32 | 1.32 | 3264.23 | 0.1012 | 1.55 | 0.1580 |
| Vertebra_C2_BM | Bone Marrow (Red) | 2471.32 | 1.32 | 3264.23 | 0.1012 | 1.55 | 0.1580 |
| Vertebra_C3_BM | Bone Marrow (Red) | 2471.32 | 1.32 | 3264.23 | 0.1012 | 1.55 | 0.1580 |
| Vertebra_C4_BM | Bone Marrow (Red) | 2471.32 | 1.32 | 3264.23 | 0.1012 | 1.55 | 0.1580 |
| Vertebra_C5_BM | Bone Marrow (Red) | 2471.32 | 1.32 | 3264.23 | 0.1012 | 1.55 | 0.1580 |
| Vertebra_C6_BM | Bone Marrow (Red) | 2471.32 | 1.32 | 3264.23 | 0.1012 | 1.55 | 0.1580 |
| Vertebra_C7_BM | Bone Marrow (Red) | 2471.32 | 1.32 | 3264.23 | 0.1012 | 1.55 | 0.1580 |
| Vertebra_T1_BM | Bone Marrow (Red) | 2471.32 | 1.32 | 3264.23 | 0.1012 | 1.55 | 0.1580 |
| Vertebra_T2_BM | Bone Marrow (Red) | 2471.32 | 1.32 | 3264.23 | 0.1012 | 1.55 | 0.1580 |
| Vertebra_C1_CB | Bone (Cortical) | 1435.18 | 2.04 | 2923.94 | 0.0203 | 3.50 | 0.0709 |
| Vertebra_C2_CB | Bone (Cortical) | 1435.18 | 2.04 | 2923.94 | 0.0203 | 3.50 | 0.0709 |
| Vertebra_C3_CB | Bone (Cortical) | 1435.18 | 2.04 | 2923.94 | 0.0203 | 3.50 | 0.0709 |
| Vertebra_C4_CB | Bone (Cortical) | 1435.18 | 2.04 | 2923.94 | 0.0203 | 3.50 | 0.0709 |
| Vertebra_C5_CB | Bone (Cortical) | 1435.18 | 2.04 | 2923.94 | 0.0203 | 3.50 | 0.0709 |
| Vertebra_C6_CB | Bone (Cortical) | 1435.18 | 2.04 | 2923.94 | 0.0203 | 3.50 | 0.0709 |
| Vertebra_C7_CB | Bone (Cortical) | 1435.18 | 2.04 | 2923.94 | 0.0203 | 3.50 | 0.0709 |
| Vertebra_T1_CB | Bone (Cortical) | 1435.18 | 2.04 | 2923.94 | 0.0203 | 3.50 | 0.0709 |
| Vertebra_T2_CB | Bone (Cortical) | 1435.18 | 2.04 | 2923.94 | 0.0203 | 3.50 | 0.0709 |
